# Serum albumin coated stellate mesoporous silica nanocomposites inhibit metastatic outgrowth in zebrafish embryos

**DOI:** 10.1101/2025.06.24.661256

**Authors:** Nandini Asokan, Vincent Mittelheisser, Arnaud Jablonski, Alexandre Adam, Joëlle Bizeau, Sébastien Harlepp, Vincent Hyenne, Olivier Lefebvre, Jacky G. Goetz, Mariana Tasso, Damien Mertz

## Abstract

Mesoporous silica-based nanoparticles (NPs) are promising tools for developing targeted therapeutic interventions in cancer. Endowed with a large pore silica shell suitable for drug encapsulation and with a responsive magnetic core, iron oxide stellate mesoporous silica (IO STMS) NPs stand out. Yet, their impact and potential toxicity on relevant *in vivo* models has not been carefully tested yet. Herein, we assessed the impact of these IO STMS nanocomposites in a syngeneic metastasis assay in zebrafish. NPs were surface-modified with human serum albumin (HSA) and loaded or not with the chemotherapeutic doxorubicin (DOX). *In vitro,* DOX-loaded NPs were expectedly more toxic to zebrafish melanoma (Zmel) cells than no-DOX NPs. In zebrafish embryos, the NPs were rapidly distributed through blood circulation and were found to colocalize over time with the vascular endothelium and local macrophages. Suprisingly, the NPs efficiently reduced the outgrowth of Zmel tumoral masses in an experimental metastasis assay in zebrafish embryos regardless of their loading with DOX. The anti-metastatic effect of these NPs was further improved by increasing the amount of HSA coating, also resulting in higher embryo survival. Altogether, IO STMS NPs showed promising cytotoxic effects on a relevant zebrafish metastasis model, inhibiting metastatic outgrowth *in vivo* independently of the drug loading. This opens the door to further testing for better exploiting their targeting and drug delivery potentialities.

## INTRODUCTION

The application of nanotechnology in cancer medicine offers the possibility to improve the efficiency of current anti-cancer therapies. In recent years, a wide variety of new nanoparticles (NPs) were designed for drug delivery, imaging and therapeutic purposes^1,2^. With successful lab-based and pre-clinical screenings for toxicity and functional efficacy, these nanoplatforms could improve the quality of life in cancer patients during treatment and along their life span. Of the myriad of NPs investigated in the past years -including soft polymer, lipid-based and inorganic nanoparticles-, it is reported that at least 80 NP-based therapies are under clinical investigation and 15 nanomedecines are currently approved^3^. However, among the main challenges to be solved, the design of nanomedecines with efficient targeting, controlled biodegradability, favorable scaling-up, or the possibility to control drug delivery with local or remote stimuli, are the principal issues to leverage.

Regarding specifically silica-based nanoparticles, to date there are mainly two theranostic nanoformulations used in clinical phases: Aurolase®, a silica core capped with a gold shell for near infrared light (NIR) treatment of prostate cancer and Cornell dots®, which are small silica dots (∼ 5 nm size) functionalised with an RGD peptide targeting ligand and a positron emission tomography (PET) probe for the detection of metastatic cancer^4^. To date, there is no mesoporous silica translation in the clinics while they have the advantage of drug transport as compared to non-porous silica. Mesoporous silica nanoparticles (MSNs), with their unique property to encapsulate small and large biomolecules (e.g. RNA, proteins), display good biocompatibility and biodegradability ensuring a promising use for drug delivery applications^5^. Various mild chemical methods allow to finely tune their pore structure, for instance by playing on the porogen surfactant lenght or on its counter ion, enabling a high versatility in drug loading or protein encapsulation^6,7^. Regarding their biodegradability, recent *in vitro* and *in vivo* (mouse) studies indicate that MSNs degrade in periods ranging from several days to several weeks and that the main factors governing degradation and clearance are the NP concentration, their degree of agglomeration and their pore size^8^. Moreover, the integration of a biocompatible inorganic core into such silica structure to afford localized treatment (heating, formation of reactive oxygen species) under external fields (light, alternating magnetic field) appears as a very appealing approach for future treatments.

Using sol-gel techniques in the presence of biocompatible inorganic cores (such as iron oxide, IO), core-shell nanomaterials have been developed, integrating the combinatorial properties of both, core and shell within a single nanostructure. At this respect, we have recently developed a new generation of core-shell iron oxide MSNs featuring a silica shell of stellate morphology (STMS) and controlled pore structure^9^. This stellate shell, formed in the presence of cetytrimethylammonium tosylate (CTATos) surfactant as a dendritic pore-directing agent, is fully tunable and exhibits a surface area of approximately 500 m².g^−1^ with a pore size ranging from 10 to 15 nm. These IO STMS NPs were efficient at generating cancer cell death *in vitro* under magnetic hyperthermia, and were shown to be potential diagnostic markers for fluorescence and magnetic resonance imaging^10^. Furthermore, when loaded with doxorubicin (DOX), these NPs behaved as effective drug carriers and as prospective photothermal agents^11^. However, to date they were never assessed in an *in vivo* model, which is the purpose of this work.

In recent years, zebrafish (*Danio Rerio*) has rapidly become an established animal model system to study NPs^12–18^. The transparency of the zebrafish embryo allows for direct visualization of NP biodistribution and of their interactions with host cells *in vivo*^19,20^. Furthermore, for a wide range of nanomaterials, recent studies indicate that NP behavior and cellular uptake process are very similar between zebrafish and mammals^21^. Noteworthy, the availability of fluorescent zebrafish reporter lines allows to investigate NP interactions with host cells, including endothelial cells and macrophages^20,22,23^. Besides, zebrafish xenograft models are very well established and have cancer avatars of different cancer cell types^24–27^. These models have been used for screening a large number of drugs and drug formulations for efficacy and toxicity^28–30^. Together with these characteristics, their low maintenance cost and amenability to high-throughput screens place the zebrafish embryo as an ideal model for screening nanobiomaterial biodistribution, toxicity, bioaccumulation, and *in vivo* fate^28–30^.

To date, the vast majority of NP assessment studies on zebrafish focuses only on toxicity^31–35^. Few studies have evaluated the toxicity of selected therapeutic MSNs in zebrafish^36–38^, their biodistribution and *in vivo* delivery potential^39–41^. NP distribution *in vivo* is highly influenced by endothelial cells, and only few studies in zebrafish have demonstrated active uptake of these NPs by endothelial cells^20,21^. Another cell type that actively uptake NPs are macrophages, but NP uptake by macrophages has been shown to be inefficient for cancer therapy as it decreases the chances of NP accumulation in tumor sites^42,43^. On the contrary, recent studies suggest that direct targeting of macrophages may be advantageous as these cells can be recruited to tumor sites^44–46^. Furthermore, among various drug formulations investigated in zebrafish for drug delivery against cancer, doxorubicin encapsulated by different nanocarriers on a zebrafish xenograft model has shown reduction in tumor burden^47–51^. Further *in vivo* analysis of the dissemination, cellular uptake, toxicity and anti-metastatic behavior of IO STMS NPs is thus warranted.

In this work, we adressed for the first time an *in vivo* study with core-shell IO STMS NPs. These NPs are well-characterized nanoplatforms that have superparamagnetic iron oxide cores and a versatile stellate mesoporous shell whose morphology and pore size can be tuned^52^. In these designs, the core-shell NPs were surface-modified with a tight human serum albumin (HSA) layer using isobutyramide (IBAM) grafts as efficient binders^53^. *In vivo*, the NP surface-coverage with HSA is vital to limit NP opsonization while in circulation^54,55^, yet whether increasing HSA coverage provides any additional benefit remained to be tested. Herein, we exploited a syngeneic zebrafish melanoma model to study the impact of HSA-coated IO STMS NPs, loaded or not with DOX, on their anti-metastatic potential. While DOX loading provided a significant advantage in cytotoxicity of Zmel cells *in vitro*, it had no impact when NPs were synchronously injected with metastatic cells *in vivo* in zebrafish embryos and assessed for their anti-metastatic potential. Yet and interestingly, we report that NPs significantly reduced metastatic outgrowth and embryo survival, and that by increasing HSA coating a further enhancement in tumor reduction and in embryo survival were revealed.

## MATERIALS AND METHODS

### Chemicals

Tetraethyl orthosilicate (TEOS, ≥99.0%)), cetyltrimethylammonium tosylate (CTATos, ≥98.0%), 2-amino-2-(hydroxymethyl)-1,3-propanediol (AHMPD, ≥99.9%), ammonium nitrate (NH4NO3), (3-amino propyl)triethoxysilane (APTES, 99%), isobutyryl chloride (IBC, 98%) and triethylamine (Et3N, ≥99%) were obtained from Sigma–Aldrich (France). Nitric acid 70% (HNO3, 70%) and *N*,*N*-Dimethylformamide (DMF, ≥99.9%) were purchased from Carlo-Erba. Iron(III) stearate (FeSt3) and fluorescein isothiocyanate (FITC) were obtained from TCI. Oleic acid (99%) was purchased from Alfa Aesar while squalane (99%) was purchased from Acros Organic. Doxorubicin hydrochloride was obtained from OChem Inc. Human serum albumin (HSA) were purchased from Sigma Life Science. Dimethyl sulfoxide (DMSO) was obtained from Roth.

### Cell culture

DMEM, non-essential amino acids and 0.05% trypsin were from Gibco. Fetal Bovine Serum (FBS), penicillin (50 units/mL) and streptomycin (50 µg/mL) were from Thermo Fisher Scientific. 1-Phenyl-2-thiourea (PTU) and Tricaine (ethyl-3-aminobenzoate-methanesulfonate) were from Sigma-Aldrich.

### Synthesis of NPs and DOX loading

IBAM-modified IO@STMS NPs (IO@STMS@IBAM) synthesis was as previously described^11^. For drug loading, doxorubicin hydrochloride (DOX) (2 mg, 3.7 µmol) was dissolved in 1 mL HEPES buffer (100 mM, pH=7.5) and incubated with 2.5 mg of IO@STMS@IBAM NPs during 24 h on a stirring wheel at room temperature, protected from light. The obtained IO@STMS@DOX NPs were collected by centrifugation (14,000 *g*, 10 min) and washed three times with 4.5 mL HEPES buffer (100 mM, pH=7.5). The DOX concentration in the supernatants (after loading and washings) was measured by UV-Vis spectrophotometry to calculate the drug loading content (DLC) in the IO@STMS@DOX NPs. For this, a calibration curve of DOX in HEPES buffer (100 mM, pH=7.5) was established by measuring the absorbance at 480 nm. Finally, a DLC of 11.3 % -meaning 113 µg.DOX mg^−1^ NPs-was obtained and used for the whole biological study.

### FITC-labelling of HSA

The procedure was reported previously by Bizeau *et al*.^56^. Briefly, a molar ratio of 2 FITC for 1 HSA protein was used to ensure efficient labelling. 5 mL HSA solution (10 mg.mL^−1^) in NaHCO_3_ buffer (0.1 mol.L^−1^, pH 8.5) was reacted with 59 µL of FITC solution (10 mg.mL^−1^ in DMSO) and stirred for 1 h. The mixture was protected from light during the whole procedure. After purification by dialysis against water (2 days, water change every 2 h), the final volume was measured to calculate the exact concentration. The HSA-FITC solution was aliquoted and stored at −20°C for further use.

### HSA-FITC coating of the NPs

Two very different coatings of HSA-FITC : HSA and High HSA, were deposited on IO@STMS@IBAM NPs, loaded or not with DOX.

#### HSA NPs

NPs@HSA-FITC (and NPs@DOX@HSA-FITC) were prepared by dispersing 10 mg of IO@STMS@IBAM (and IO@STMS@IBAM@DOX) NPs in 2.5 mL of a solution of HSA-FITC (0.4 mg.mL^−1^ in HEPES 100 mM, pH 7.5) resulting in a 10 wt% HSA-FITC feed weight ratio. The coating was deposited by stirring the suspensions on a rotating wheel for 1 h, at room temperature and protected from light. The NPs@HSA-FITC (and NPs@DOX@HSA-FITC) were centrifuged (12,000 *g*, 12 min) and washed twice (12,000 *g*, 12 min) with HEPES buffer (100 mM, pH 7.5). HSA NPs, with and without DOX, are thereby modified with a 10 wt% ratio of HSA. These compounds are refered as HSA NPs or HSA-Dox_NPs in the text.

#### High HSA NPs

For NPs@HighHSA-FITC, an additional incubation with 1 mL HSA-FITC at 10 mg.mL^−1^ (100% feed weight ratio) was performed to increase the protein surface coverage. For NPs@DOX@HighHSA-FITC, additional successive incubations with 1 mL HSA-FITC at 1 and 10 mg.mL^−1^ (respectively 10 and 100 % feed weight ratio) were carried out to increase the protein surface coverage. High HSA NPs are therefore modified with a 110 wt% ratio of HSA while DOX-containing High HSA NPs have a slightly higher exposure to HSA at a 120 wt%. As above, after HSA modification, the NPs were centrifuged (12,000 *g*, 12 min) and washed twice (12,000 *g*, 12 min) with HEPES buffer (100 mM, pH 7.5). These compounds are refered as High HSA-NPs or High HSA-Dox_NPs in the text.

Both, HSA and High HSA surface contents were calculated from the measurement of the fluorescence emission intensity of the supernatants at 519 nm (excitation at 490 nm). The HSA-coated NPs (with and without DOX) were stored at 4°C in 100 mM HEPES buffer, pH 7.5 at a high concentration (i.e. in the mg.mL-1 range) prior to their use in the biological assays.

### Physicochemical characterization of the NPs

Once the IO@STMS NPs synthesized and the CTATos surfactant extracted from the pores, the NPs were imaged with a JEOL 2100 TEM instrument operating at 200 kV. Image J was used to analyse the size distribution of the NPs. IBAM graft density was evaluated using thermogravimetric analysis (TGA) which was performed on a STD Q600 (TA Instruments, 25-800°C, a heating rate of 5°C/min, under air flow rate of 50 mL.min^−1^). Hydrodynamic size distributions of the NPs with and without DOX, and at HSA and High HSA ratios, were evaluated by dynamic light scattering (DLS) in the Intensity mode using a Malvern Nanosizer instrument operating at ambient temperature. Colloidal stability was evaluated in different buffers: HEPES (100 mM, pH 7.5), PBS (pH 7.4), and cell culture medium (RPMI).

### Cell culture

Zebrafish melanoma cells (Zmel dark and Zmel-tdTomato) were cultured in DMEM containing 10% Fetal Bovine Serum (FBS), 1% penicillin (50 units.mL-1) and streptomycin (50 µg.mL-1) and 1% non-essential amino acids, and were maintained at 28°C in a humidified incubator supplied with 5% CO_2_. Cells were harvested when reaching 85% confluence. Cells were detached with 0.05% trypsin, treated for 3 min at 28°C.

### Cell viability

For cytotoxicity experiments, 125,000 Zmel dark or Zmel-tdTomato were plated in a 24-well plate and let overnight. The day after, medium was changed for medium containing increasing concentrations (0 to 200 µg.mL-1) of NPs, doxorubicin-loaded NPs (Dox_NPs) or matching concentrations of doxorubicin.HCl (DOX). The NPs were surface-coated with HSA-FITC, as previously described. After 48h, cells were washed, fixed with PFA 4% and stained for 1h with 2% crystal violet. Excess crystal violet was washed with tap water and plates were let dry before imaging with an iBright 1500 imaging system. Images were loaded on ImageJ and crystal violet mean grey intensity in each well was measured after oval selection definition.

### Cellular internalization *in vitro*

For internalization experiments, 500,000 Zmel-tdTomato cells were plated in a 4-well LabTek chamber with coverlid and let overnight. The day after, medium was changed for medium containing 100 µg.mL-1 NPs or Dox_NPs in complete medium. The NPs were surface-coated with HSA-FITC, as previously described. After 5h of incubation at 28°C, cells were washed and fixed with PFA 4% before mounting in Fluoromount/DAPI. Slides were imaged using a LSM800 Zeiss confocal microscope with a Plan Apo ×40 oil (NA 1.4) objective. The excitation wavelenghts were: 405 nm for DAPI; 488 nm for FITC, and 554 nm for tdTomato.

### Intravascular injection of NPs in zebrafish embryos

Zebrafish embryos were obtained from the following strains: Wildtype-AB, Tg(Kdrl.hsa-HRAS:mCherry)^57^, Tg(mpeg1:Gal4UAS:NTR-mCherry)^58,59^. Embryos were maintained at 28°C in Danieau 0.3X medium, supplemented with 1-Phenyl-2-thiourea after 24 h post fertilization (hpf). All injection experiments were carried out at 48 hpf and imaged between 48 hpf and 72 hpf. All animal procedures were performed in accordance with French and European Union animal welfare guidelines and supervised by local ethics committee (Animal facility #A6748233; APAFIS #2018092515234191). At 48 hpf, zebrafish embryos were dechorionated and mounted on a 0.8% low melting agarose pad containing 650 µM tricaine (ethyl-3-aminobenzoate-methanesulfonate). Embryos were injected intravascularly in the duct of Cuvier with 2 x 13.9 nL of NPs (with and without doxorubicin, at 0.5 mg/mL) with a Nanoject microinjector 2 (Drummond) under a M205 FA stereomicroscope (Leica), as described previously^60,61^. The NPs were surface-coated with HSA-FITC, as detailed above. At 3 hpi (hours post-injection) and 24 hpi injected embryos were imaged using the Olympus IXplore Spin inverted spinning-disk microscope at 30x with a 1.05 NA (silicone) objective focusing on the head and caudal hematopoietic tissue region (CHT) (488 nm laser at 2% for 100 ms / 561 nm laser at 15% for 300 ms). Quantification of NP colocalization with endothelial cells and macrophages was done using an in-house macro in imageJ (Fiji). Embryo survival after NP administration was monitored every day until 4 dpi (days post-injection), with embryos screened for any phenotypic abnormalities. Percentage of embryo survival was plotted using a Kaplan-Meier curve. Pacemaker activity was measured by recording the heartbeat of the embryos as short time-lapses with a Stereomicroscope (Leica M205 FA). Heartbeats were manually counted.

### Assessment of NP effect onto macrophage number and length of main vessels

Quantification of macrophage number in the head and trunk regions was performed on processed single Z planes based on intensity thresholding using Cell Profiler. The Intersomitic vessel (ISV) length was measured from the point where the ISV sprouts from the dorsal aorta up to the tip of the ISV using imageJ. Similarly, Mesencephalic vein (MsV) length was measured from the point where the MsV extends from the dorsal side up to the tip of the MsV reaching artery using ImageJ.

### Tumor co-localization assay *in vivo*

This experiment was carried out with two types of NPs: NPs with HSA and High HSA surface content, both with and without encapsulated doxorubicin. In a typical experiment, about 200 zebrafish melanoma cells (Zmel-tdTomato = Zmel_tdT) were injected into the Duct of Cuvier at 36 hpf on a wildtype AB embryo. Embryos properly grafted with zebrafish melanoma cells were divided into 3 groups. At 48 hpf, 2 x 13.9 nL of PBS, NPs or Dox_NPs at 0.5 mg.mL-1 were injected into the circulation of the larvae of the pre-divided groups. The NPs were surface-coated with HSA-FITC, as previously described. At 3 hpi, 24 hpi and 4 dpi, injected embryos were imaged using spinning disk microscopy focusing on the caudal hematopoietic tissue region (CHT). Tumor volume at different time points was quantified with the Imaris (Interactive microscopy image analysis) software. Data are normalized according to the initial volume at 3 hpi for each embryo. Embryo survival was quantified at the defined time points.

### Colocalization analysis

Colocalization was analyzed on processed single Z planes (Despeckle and Gaussian blur filter in Fiji) based on intensity thresholding using Cell Profiler. Areas of overlapping regions between NPs and tumor cells were quantified. The percentage of NP colocalization with the tumor cell was calculated as a ratio of the NP overlap area to the total area of NP objects times 100.

### Statistical analysis

Statistical analysis was done using GraphPad Prism (version 10.2). Normality of the data was confirmed using Shapiro-Wilkson test. According to the results, different statistical tests were used, as described in each specific section and in figure legends. For data that follow a Gaussian distribution, an unpaired t test was used; if not, a Mann-Whitney approach was applied. Two-way Anova analysis was applied to analyzed tumor growth over time using the two-stage linear step-up post-test procedure of Benjamini, Krieger and Yekutiel. Survival was analyzed using the Log-rank (Mantel-Cox) test with a Bonferroni post-test setting the number of comparisons, k, to 3. Violin plots are employed to present the data, with median and quartiles included in the plots. p-values smaller than 0.05 were considered as statistically significant. * = p<0.05, ** = p<0.01, *** = p<0.001, **** = p<0.0001.

## RESULTS AND DISCUSSION

### Synthesis and characterization of the HSA-coated NPs

The NPs surface-modified with human serum albumin (HSA), denoted HSA NPs, and loaded or not with doxorubicin (DOX) molecules, were synthesized as previously reported by Adam *et al*.^11^ (see **Scheme S1** for the whole chemical strategy). This previous work from our group focused on the evaluation of the photothermal properties of the core-shell NPs, and on their drug loading/release capabilities as a function of pH and surface chemistry. Here, the aim is to analyze this well-characterized and versatile nanosystem in an *in vivo* condition, under blood circulation, and upon interaction with circulating tumor cells, macrophages and endothelial cells present in the vasculature. Briefly, the iron oxide (IO) cores were obtained by thermal decomposition, a method allowing a precise control over nanoparticle size and composition. Transmission electron microscopy (TEM) images indicate monodisperse spherical NPs with an average size distribution of 26.4 ± 3.4 nm, while X-ray diffraction (XRD) confirms the spinel phase structure (Fe_3-x_O_4_, **Figure S1.A-C**). The stellate mesoporous silica (STMS) shell was grown around the iron oxide cores using the sol-gel procedure in the presence of CTATos as the templating agent directing the stellate large pore morphology. Then, the surfactant was removed to free the pores and avoid any further release in biological media. TEM images confirm the synthesis of individual core-shell IO@STMS NPs having an average diameter of 119.7 ± 12 nm and optimal colloidal stability in ethanol, as determined by dynamic light scattering (DLS) (**Figure 1.A-C**). Next, the surface was chemically modified in two steps with isobutyramide (IBAM) moieties covalently grafted at the extremities of condensed aminosiloxane groups (**Fig. S1.D,** Thermogravimetric analysis). IBAM groups were previously reported as suitable binders for DOX loading and HSA immobilization^11^. DOX loading was set here at one specific loading condition ensuring a drug content of 113 µg DOX.mg-1 of NP for the whole biological study. DOX-loaded NPs were afterwards exposed to a human serum albumin protein (HSA) solution at a 10 wt% ratio relative to the NP mass, giving rise to an HSA content on the NP surface of 96 µg HSA.mg-1 of NP. This weigh ratio ensured an excellent yield (close to 100% impregnation) and the possibility of DOX release at pH mimicking intracellular conditions (∼5.5) with a great efficiency^11^. Further, an HSA coating in the range 100-200 µg HSA.mg-1 ensured HSA NP interactions with HeLa cancer cells with suitable cell uptake (at NP concentrations in cell culture <30 µg.mL-1)^10^. DOX and HSA contents were calculated from supernatant analysis by UV-Vis spectrophotometry and fluorescence spectroscopy, respectively, using calibration curves in HEPES buffer (**Figures S1.E,F**). Next, the colloidal stability of the NPs was evaluated in different biological media. Dynamic light scattering (DLS) measurements in RPMI cell culture medium provided average HSA NP diameters of 234 ± 4 nm (DLS Z-average value), with good colloidal stability. On the other hand, NPs tended to form aggregates of micron-size in buffers such as HEPES or PBS. For DOX loaded NPs, results are presented in **Figure 1.D-F** while the whole study with and without DOX is presented in **Figure S2** with the same conclusions.

**Figure 1.**
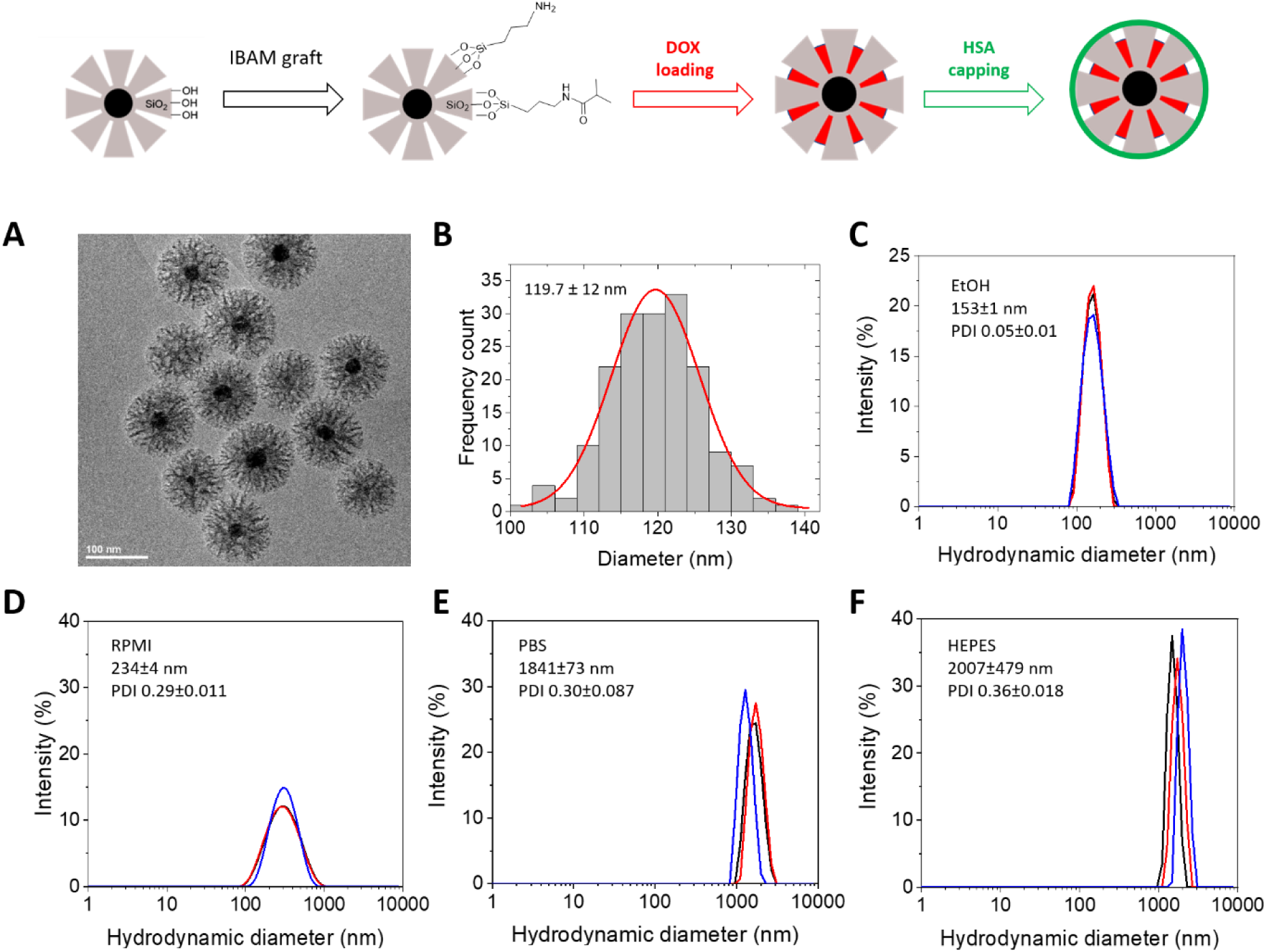
Main physicochemical features of the core-shell NPs surface-modified with HSA. Top: Summary of the chemical steps, with initial silica shell modification with IBAM functional units, followed by doxorubicin (DOX) loading and final HSA surface immobilization. A) TEM image of the core-shell NPs formed by an iron oxide core and an stellate mesoporous silica shell. B) Associated NP size histogram. C) NP hydrodynamic size distribution in the storage solvent ethanol prior DOX loading and HSA modification (DLS measurement). D-E-F) NP hydrodynamic size distribution in three different biological media (RPMI cell culture medium, PBS, and HEPES) for the core-shell NPs loaded with DOX and having HSA coating (DLS measurement). DLS runs were done in triplicate in each medium as indicated in the DLS graphs. The average hydrodynamic diameter of the NP distribution is indicated as inset. DLS = Dynamic Light Scattering. PDI = PolyDispersity Index.

### Drug-loaded HSA NPs are cytotoxic to zebrafish melanoma cells *in vitro* but not HSA NPs

We first assessed the impact of these engineered NPs on a relevant metastatic melanoma model (Zmel) that is compatible with the syngeneic experimental metastasis assays to be implemented further. For that, we evaluated the viability of Zmel cells (engineered to express the fluorescent TdTomato protein, named Zmel_tdT) upon 48 hours exposure to various concentrations of the HSA NPs, loaded or not with DOX (Dox_NPs), and of the drug DOX alone. Toxicity of Dox_NPs with regards to Zmel cells mirrored the one obtained with free DOX and scaled with increasing concentrations (**Figure 2.A**). Of note, high concentrations (above 50 µg.mL-1) of drug-free NPs led to mild Zmel cells toxicity. When assessing the interactions of NPs with Zmel cells at the chosen NP concentration of 100 µg.mL-1, we observed micron-scale clusters at the cell surface independently of DOX loading (**Figure 2.B**). Altogether, this suggests that highly-concentrated NPs (above 50 µg.mL-1) are detrimental to Zmel tumor cells and that DOX loading provides a concentration-dependent anti-tumoral benefitial effect. Noteworthy, cell toxicity was distinctly higher for Dox_NPs than for the NPs without the drug, clearly highlighting a drug-specific cellular response.

**Figure 2.**
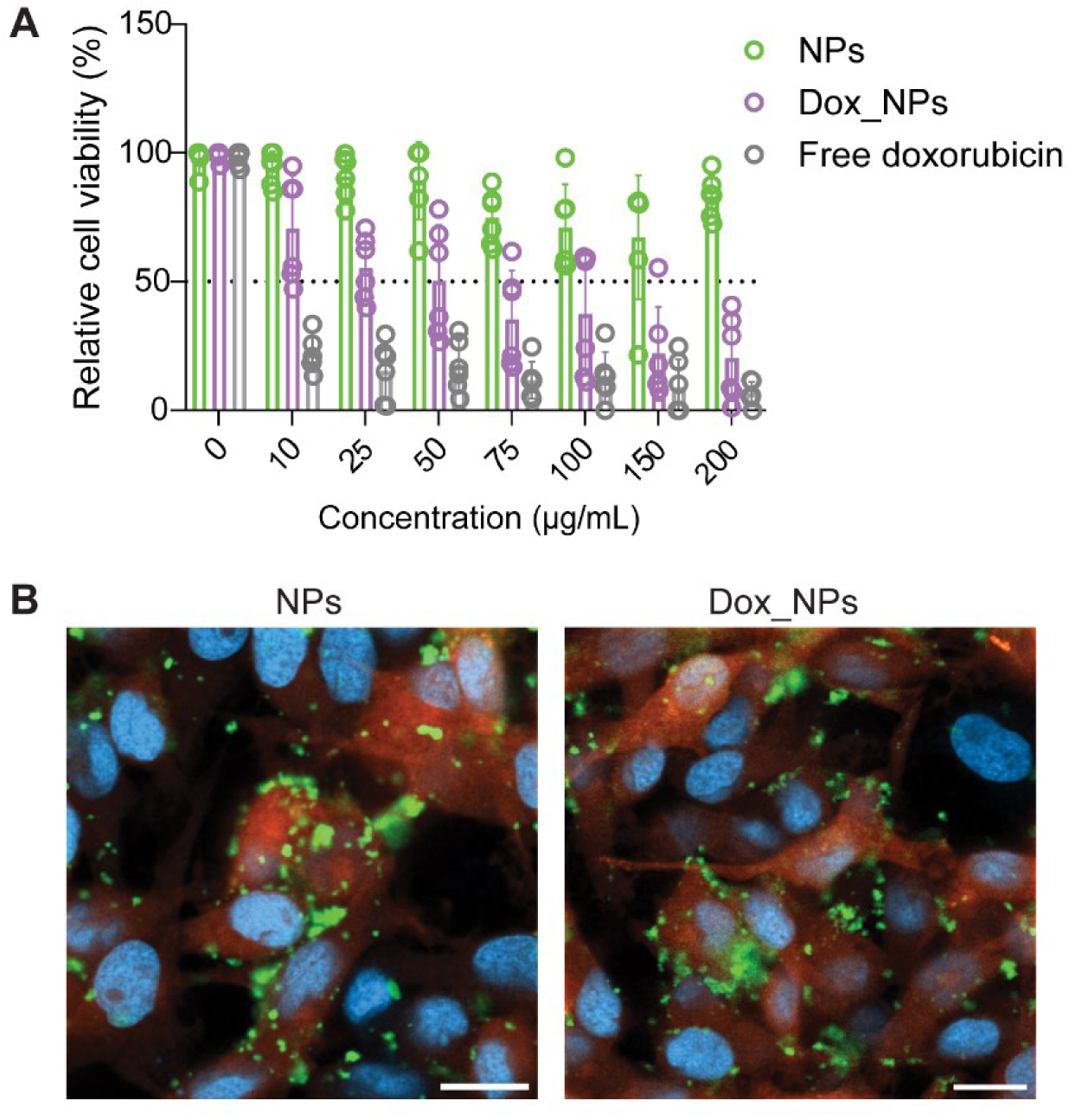
HSA NPs affect zebrafish melanoma cell viability *in vitro*. A) Quantification of zebrafish melanoma cell (Zmel_tdT) viability upon a 48 hour treatment with HSA NPs (with and without doxorubicin) using crystal violet assay. Free doxorubicin (DOX) in solution was also included as positive control. Results are normalized to untreated cells. DOX-containing NPs (Dox_NPs) showed a dose dependent effect similar to that of free DOX while cell viability is only affected at high NP concentrations for the NPs without the drug. Data are displayed as mean ± SD. B) Representative images of NP (in green) interactions with the zebrafish melanoma (Zmel_tdT) cell line (in red). Cells were exposed during 5 hours at 28°C to 100 µg.mL-1 NPs or doxorubicin-loaded NPs, both surface-modified with HSA-FITC. DAPI stained nuclei in blue. Scale bar = 10 µm.

### HSA NPs interact with the endothelium and macrophages

Probing the *in vivo* behavior of NPs is instrumental to study their interactions with the host or their circulation time, which ultimately determines the efficacy of the therapeutics. Because the size, shape, and surface features of NPs can be tuned, *in vivo* tracking is the only method that enables a clear analysis of how these parameters control their biodistribution, preferential accumulation in specific organs, toxicity, and final fate^62–64^ In that context, the zebrafish embryo has proven to be a model of choice for NP toxicity, biodistribution and drug targeting efficacy^65–67^.

Serum albumin nanoparticles have been extensively studied as anti-cancer drug delivery agents, due to their efficient uptake by cancer cells^68–70^. Upon uptake, the lysosomal degradation of serum albumin NPs induces cancer cell death through the release of their encapsulated drugs^71^. Among FDA approved drugs, DOX is known for its selective tumor localization and pharmacokinetic properties, and it has been encapsulated inside nanocarriers^72–74^. Although zebrafish do not have serum albumin proteins, it has been shown that HSA can be processed in the liver of juvenile zebrafish indicating the relevance of the model^75^. Moreover, the size of the HSA-modified NPs did not allow for direct absorption of NPs from fish water. Hence, microinjection of NPs into the blood stream was preferred and it enabled NP dissemination throughout the vasculature, in agreement with the biodistribution of different nanoformulations injected intravascularly.

Hence, in a first *in vivo* study, we assessed the stability and biodistribution of FITC-labeled HSA NPs (with and without DOX) in a relevant animal model and in real time. When microinjected into the duct of Cuvier of transgenic zebrafish strains Tg(Kdrl.hsa-HRAS:mCherry)^57^ and Tg(mpeg1:Gal4UAS:NTR-mCherry)^58,59^ that label endothelial cells and macrophages, respectively (**Figure 3.A,D**), the HSA-FITC NPs disseminated throughout the embryo irrespective of the transgenic line. Regarding their interactions with the endothelial system, many NPs and Dox_NPs were found in the caudal plexus area and in the head, close to optic and other brain vessels. Further, the NPs without the drug remained much longer in circulation than the Dox_NPs (found static at 3 hpi already, **Figure 3.B**), suggesting that the presence of DOX has an effect on the fate of these NPs in the zebrafish blood stream. Both NPs and Dox_NPs showed similar percentage of colocalization with endothelial cells (**Figure 3.C**) at 3 hpi, indicating that DOX encapsulation had no distinctive effect on this phenomena. Confocal imaging confirmed that some NPs were internalized by endothelial cells independently of the DOX loading. Interestingly, we noticed that Dox_NPs were more prone to spread across the extravascular space at 24 hpi (**Figure S3.A,B**). Dox_NPs rapidly form immobile aggregates in proximal yet extravascular regions.This suggests that Dox_NPs might interact with specific host cell types or induce cellular mechanisms that can aid their extravasation. In general, NPs are unable to escape interactions with other host proteins and the consequent formation of a protein corona. This phenomenon ends up enabling their visibility to phagocytic cells.

**Figure 3.**
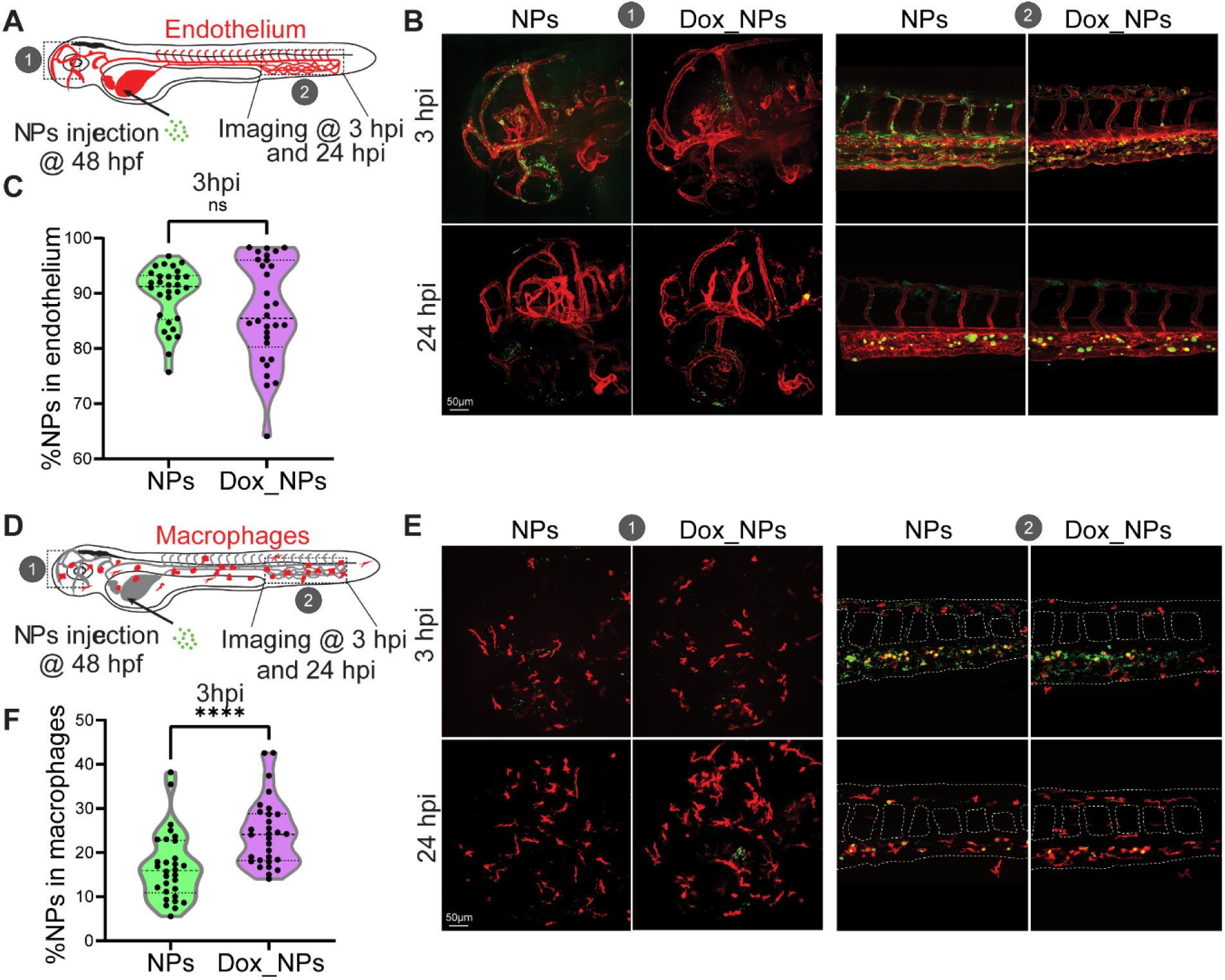
Biodistribution of HSA NPs and their interaction with endothelial cells and macrophages. A) Description of the experimental setup: 48 hours post-fertilization (hpf), Tg(Kdrl.hsa-HRAS:mCherry) zebrafish embryos (endothelial cells in red) are injected intravascularly *via* the duct of Cuvier with both, NPs and DOX-containing NPs (Dox_NPs) surface-modified with HSA-FITC. Embryos are imaged in the head (1) and caudal plexus area (2) 3 or 24 hours post injections (hpi). B, C) Representative images (B) of the NPs in the head (1) and caudal plexus (2) areas, and quantification of NPs colocalizing with the endothelium (C). At 3 hpi, there is no difference between the NPs and Dox_NPs injected groups (Mann-Whitney test, 3 independent experiments, 30 embryos in total). D) Description of the experimental setup: 48 hours post-fertilization (hpf), Tg(mpeg-ntr:mCherry) zebrafish embryos (macrophages in red) are injected intravascularly *via* the duct of Cuvier with both NPs and DOX-containing NPs (Dox_NPs) surface-modified with HSA-FITC. Embryos are imaged in the head (1) and caudal plexus area (2) 3 or 24 hpi. E, F) Representative images (E) of the NPs in the head (1) and caudal plexus (2) areas, and quantification of NPs colocalizing with macrophages (F). At 3 hpi, Dox_NPs displayed a significantly higher colocalization with macrophages as compared to the NPs without the drug (Mann-Whitney test, p<0.0001, 3 independent experiments, 30 embryos in total). Scale bar 50µm.

Interestingly, caudal plexus area homes the origin of macrophages^76^; it might thereby be possible that high accumulation of NPs in this area is due to macrophage engulfment. To confirm this hypothesis, NPs were microinjected into the zebrafish fluorescently labeled macrophage reporter line Tg(mpeg1:Gal4UAS:NTR-mCherry)^58,59^. Upon injection, the NPs and Dox_NPs were found distributed throughout the embryo (**Figure 3.E**). At 3 hpi, Dox_NPs were found colocalizing with macrophages at a higher percentage than their counterparts without DOX (**Figure 3.F),** pointing at a potential faster clearance of Dox_NPs by macrophages. Additionally, at a later time point (24 hpi), around 35 % of both NPs, with and without DOX, colocalized with macrophages (**Figure S3.C**), indicating that their removal from circulation at later stages in the zebrafish is similar, irrespective of the encapsulated drug. In this report, we did not address the uptake of NPs by neutrophils, as it has been shown that neutrophils are less involved than macrophages in internalizing NPs and other vesicles when administered^23,77,78^.

Hence, overall, by analyzing the colocalization between the NPs, the macrophages and the endothelium, the here-proposed method efficiently detects most NPs injected into the system and enables the insightful quantification of NP fractions present in colocalizing regions. The elevated colocalization with macrophages, specially at short-times after injection, can be associated with their phagocytic activity, though this remains to be fully elucidated.

### HSA NPs do not affect zebrafish vascular and innate immune embryo physiology

We further analyzed endothelial and macrophage homeostasis upon injection of NPs in 48 hpf Tg(Kdrl.hsa-HRAS:mCherry)^57^ and Tg(mpeg1:Gal4UAS:NTR-mCherry)^58,59^ zebrafish embryos (**Figure 4.A,D**). We used the length of the mesencephalic vein (MsV) and of the intersomitic vessel (ISV) as a proxy for vascular homeostasis. At 24 hpi, the vessel anatomies in the head and trunk of the injected embryos had similar lengths irrespective of the injected NP (with and without DOX, **Figure 4.B,C**). Similarly, at 24 hpi the ramified shape of macrophages was maintained upon microinjection of the NPs (**Figure 4.E**), suggesting there was no activation of macrophages at this time point, which can be assessed by their round shape. Hence macrophage homeostasis at 24 hpi remained unperturbed in both NPs and Dox_NPs injected zebrafish. Further, the number of macrophages in the head and trunk was also maintained irrespective of their injection with NPs and Dox_NPs (**Figure 4.F**), and pacemaker activity was not altered (**Figure S4.A**). For the latter, the unperturbed heart rate in the presence of DOX is a noteworthy result given the known cardiotoxicity^79,80^ of this chemotherapeutic drug.

**Figure 4.**
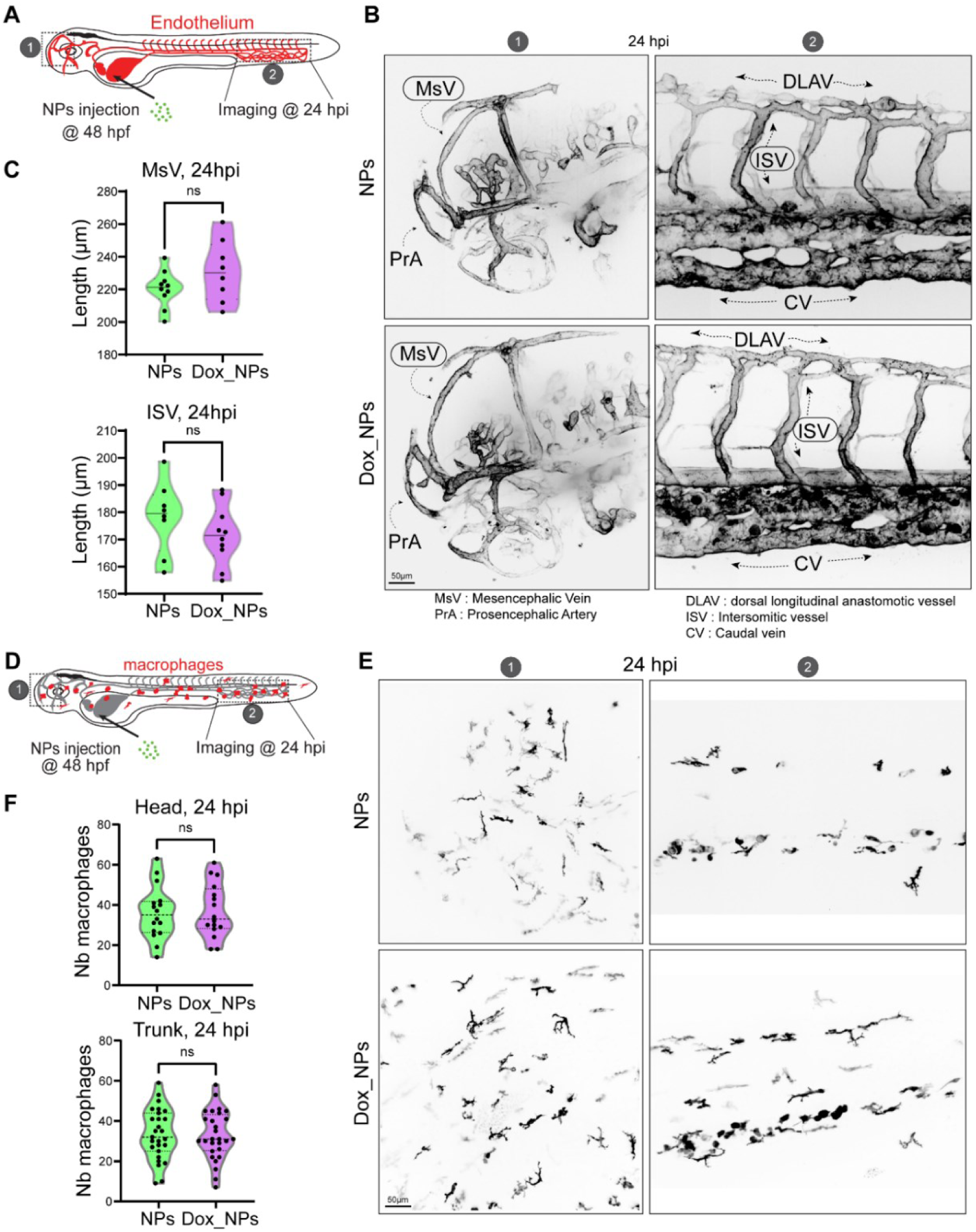
Endothelial and macrophage homeostasis remain unperturbed. A) Description of the experimental setup: 48 hours post-fertilization (hpf), Tg(Kdrl.hsa-HRAS:mCherry) zebrafish embryos (endothelial cells in red) are injected intravascularly *via* the duct of Cuvier with both NPs and DOX-containing NPs (Dox_NPs) surface-modified with HSA-FITC. Embryos are imaged in the head (1) and caudal plexus area (2) at 24 hpi. B) Representative images of Mesencephalic vein (MsV) in the head (1) and Intersomitic vessel (ISV) in the caudal plexus (2) at 24 hpi. C) Quantification of MsV (upper) and ISV (lower) length showed no difference between the NPs and Dox_NPs injected groups at 24 hpi (Unpaired t test, 3 experiment, 8 embryos for NPs, 10 for Dox_NPs). D) Description of the experimental setup: 48 hours post-fertilization (hpf), Tg(mpeg-ntr:mCherry) zebrafish embryos (macrophages in red) are injected intravascularly *via* the duct of Cuvier with both NPs and DOX-containing NPs (Dox_NPs) surface-modified with HSA-FITC. Embryos are imaged in the head (1) and caudal plexus area (2) at 24 hpi. E) Representative images of macrophages in the head (1) and caudal plexus area (2) at 24 hpi. F) Quantification of the number of macrophages at 24 hpi showed no difference between the NPs and Dox_NPs groups in both the head (upper) and trunk (lower) areas (Unpaired t test, 3 independent experiments, 16 embryos per treatment in total for the head, 27 embryos per treatment in total for the trunk). Scale bar is 50 µm.

Altogether, these data suggest that the HSA NPs, after 24 hours of injection in the zebrafish blood circulation, do not alter the main morphological features of the embryo (length of main vessels, number of macrophages, absence of noticeable phenotypic modifications).

### HSA NPs reduce tumor growth *in vivo* but affect survival

The zebrafish model, with its high similarity in genome with humans^81–83^ (around 70%), easy manipulation and amenability for high-throughput screens has been utilized as an effective xenotransplantation model without risk of rejection for almost the past two decades^24,26,30,84^. Several embryonic zebrafish xenograft studies have used doxorubicin as a loaded drug in different nanoparticle formulations and have shown reduction in tumor burden^51,73,85,86^. Albeit the promising results, there are still limitations in their clinical translation partly due to their anatomical differences from mammals and due to the lack of adaptive immune system at this developmental stage. With this in mind, we envisioned to test the HSA NPs in a zebrafish embryo model that used syngeneic zebrafish melanoma (Zmel) cells for transplantation. This cell line forms mets within 3 days after their injection in zebrafish embryos. We next tested whether HSA NPs could successfully impair metastatic outgrowth in this syngeneic experimental metastasis model. First, zebrafish wildtype embryos were intravascularly injected with fluorescently-labeled zebrafish melanoma cancer cells (Zmel_tdT) at 36 hpf. This injection was followed by injections of fluorescent HSA-FITC NPs (with and without DOX) at 48 hpf (**Figure 5.A**). Injected embryos were longitudinally imaged and assessed at 3, 24 and 96 hpi to document metastatic outgrowth (**Figure 5.B**). Both, NPs and Dox_NPs reduced metastatic outgrowth of Zmel cells (50% and 31%, respectively) (**Figure 5.B,C**) in comparison to the control PBS group. Unexpectedly, we observed a significant mortality of embryos 4 days post-injection, which reached 47 and 80 % for NPs and Dox_NPs, respectively (**Figure 5.D**). Thus, despite showing remarkable anti-metastatic potency, yet independently of DOX loading, HSA NPs displayed an unexpected mortality that warrants further investigation. As for any kind of chemotherapeutic, the NP dose, their unwanted interactions with other organs and tissues, and their removal pathways need to be fully screened and optimized.

**Figure 5.**
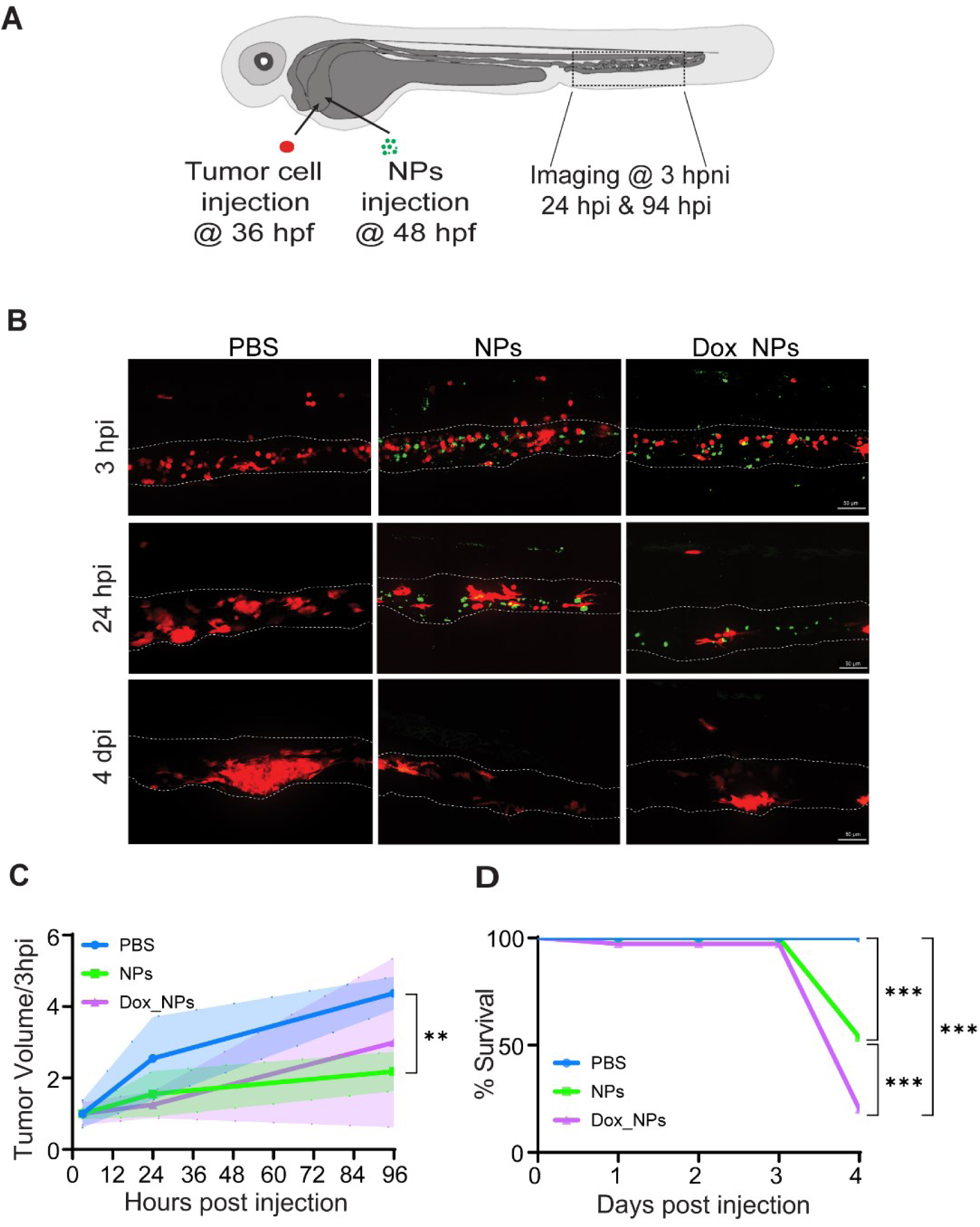
Reduction of tumor growth upon HSA NP injection. A) Description of the experimental setup: 36 hours post-fertilization (hpf), wildtype zebrafish embryos are injected intravascularly *via* the duct of Cuvier with zebrafish melanoma cells (Zmel_tdT, in red). At 48 hpf, embryos with grafted Zmel_tdT cells were divided into three groups and injected with PBS, NPs and DOX-containing NPs (Dox_NPs), respectively. The NPs (in green) are surface-modified with HSA-FITC. The caudal plexus regions were imaged at 3h, 24h and 96h post NP injection (hpi) to follow tumor growth. B) Representative images of the caudal plexus area from engrafted embryos at 3h, 24h and 96h post PBS, NPs and Dox_NPs injection. Scale bar is 50 µm. C) Quantification of tumor volume using Imaris showed a substantial tumor volume reduction with NPs and Dox_NPs at 96 hpi when compared to the PBS injected group. Data are normalized according to the initial volume at 3 hpi for each embryo (Two-way Anova followed by the two-stage linear step-up procedure of Benjamini, Krieger and Yekutieli post-test, with PBS/NPs p=0.0083, PBS/Dox_NPs p=0.081, NPs/Dox_NPs p=0.2, 1 experiment, 3 embryos/group). D) Survival analysis of tumor grafted embryos after intravascular injections of PBS, NPs and Dox_NPs showed significant embryo mortality, indicating that NPs are harmful to tumor-grafted embryos after day 3 post-injection (Log-rank (Mantel-Cox) test with Bonferroni post-test correction; PBS/NPs p=0,0003, PBS/Dox_NPs p=0,0003, NPs/Dox_NPs p=0.0069, 3 independent experiments, 27, 28 and 29 embryos in total for PBS, NPs and Dox_NPs, respectively).

Noteworthy, the *in vitro* NP experiments with Zmel melanoma cells had already underlined the fact that HSA NPs, with and without the drug, display strong unspecific interactions with the cell membrane, resulting in NP aggregates. Indeed, for the HSA NPs, 10-20% colocalization between tumor cells and NPs was obtained at all evaluated time points (**Figure S4.B**), confirming potential cell-nanomaterial interactions also *in vivo*. Herein, we assume that these non-specific interactions could be behind the underlying mechanisms that inhibit or perturb cell-cell communication, a *sine qua non* condition for tumor volume progression in this metastatic model. As it has been reported in multicellular tumor spheroids, the presence of NPs during the process of spheroid formation can alter spheroid morphology, resulting in the generation of smaller or less compact multicellular masses^87^

### High HSA-NPs further reduce metastatic outgrowth and improve survival *in vivo*

HSA is a protein that was reported to reduce the opsonization of several types of NPs in cancer models, thereby increasing their circulation time^88,89^, and to mediate the internalization of drug payloads into tumors *via* its interaction with several receptors overexpressed in tumor cells^90,91^. With this in mind, we explored the effect of increasing the HSA NP surface coverage on their anti-metastatic potential. We increased HSA immobilized on the NP surface by adding an additional HSA coating step with a 10 times more concentrated HSA protein solution as compared to the first step. This increased HSA surface concentration from 96 (previous HSA coating) to 428 µg HSA.mg-1 of NP (High HSA NP coating), resulting in 4 times more immobilized HSA at the surface of the NPs **(Figure 6.A).** This effect was attributed to the IBAM grafts at the STMS surface which allow to tighly immobilize HSA with increasing immobilized amounts as a function of the initial HSA concentrations, as previously shown by Bizeau *et al*. ^56^. We further assessed colloidal stability in different buffers: RPMI, PBS and HEPES by DLS (**Figure 6.B-E, and Figure S5.A,B**). The hydrodynamic diameter in RPMI cell culture medium of the High HSA NPs with or without DOX remained similar (and even slightly better) to the corresponding HSA NPs, suggesting overall a good colloidal stability (187 ± 8 nm for High HSA NPs versus 170 ± 7 nm for High HSA-DOX NPs) but a micron size aggregation state in PBS or HEPES.

**Figure 6.**
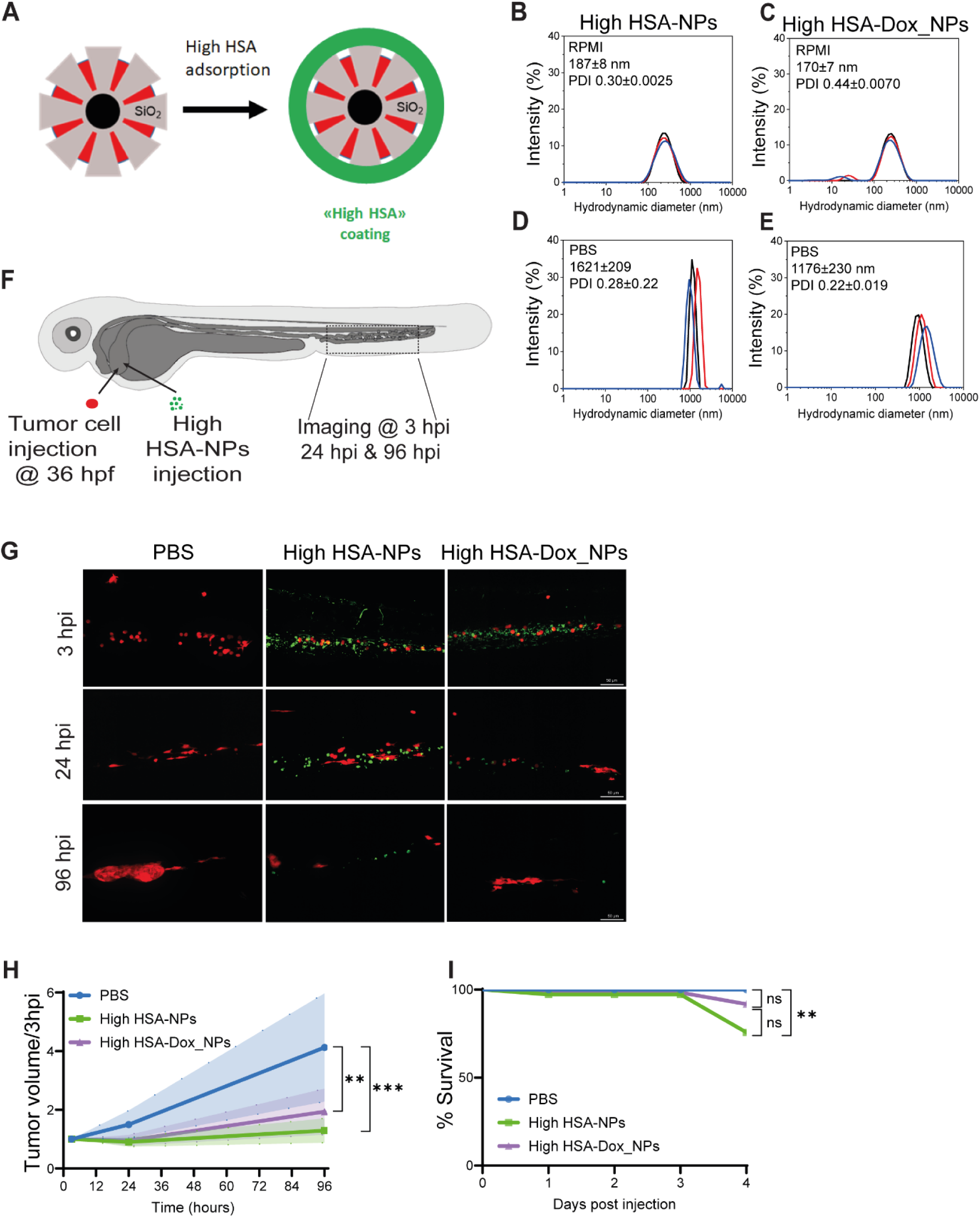
Main physicochemical features of the core-shell NPs surface-modified with High HSA and their beneficial effects onto tumor reduction and reduced mortality. A) Schematic view of the HSA adsorption onto DOX-loaded NPs for High HSA NPs. As for the previous HSA NPs, the protein is immobilized on the surface after DOX loading on the mesoporous silica shell. For High HSA NPs, protein surface coverage is increased 4-fold. Right: Dynamic Light Scattering of the High HSA NPs in RPMI (B,C) and PBS (D,E). DLS measurements were done in triplicate in each medium, as indicated in the DLS graphs. The average hydrodynamic diameter of the NP distribution is indicated as inset. F) Description of the experimental setup: 36 hours post-fertilization (hpf), wildtype zebrafish embryos are injected intravascularly *via* the duct of Cuvier with zebrafish melanoma cells (Zmel_tdT, in red). At 48 hpf, embryos with grafted Zmel_tdT cells were divided into three groups and injected with PBS, High HSA NPs and High HSA-Dox NPs. The NPs (in green) are surface-modified with High HSA-FITC. The caudal plexus regions were imaged at 3h, 24h and 96h post NP injection (hpi) to follow tumor growth. G) Representative images of the caudal plexus area from engrafted embryos at 3h, 24h and 96h post PBS, High HSA NPs and High HSA-Dox NPs injection. H) Quantification of tumor volume using Imaris showed a significant reduction of tumor volume with High HSA NPs and High HSA-Dox NPs at 96 hpi when compared to the PBS injected group. Data are normalized according to the initial volume at 3 hpi for each embryo (Two-way Anova followed by the two-stage linear step-up procedure of Benjamini, Krieger and Yekutieli post-test, with PBS/NPs p= 0.0004, PBS/Dox_NPs p= 0.0013, NPs/Dox_NPs p=0.115, 1 experiment, 3 embryos/group). I) Survival analysis of tumor grafted embryos after intravascular injection of PBS, High HSA NPs and High HSA-Dox NPs showed improved survival efficacy compared to previous HSA NPs (Log-rank (Mantel-Cox) test with with Bonferroni post-test correction, PBS/NPs p=0.003, PBS/ Dox_NPs p=0.125, NPs/Dox_NPs p=0.324, p= 3 independent experiments, 31, 30 and 32 embryos in total for PBS, NPs and Dox_NPs, respectively).

These High HSA NPs were administered to embryos as done for the HSA NPs. High HSA NPs showed similar bio-distribution *in vivo* as HSA NPs and had no impact on the overall physiology of zebrafish embryos. When subjected to experimental metastasis, we again documented a significant reduction of metastatic outgrowth at 96 hpi for both NP types, with 69% and 53% tumor volume reduction for NPs and Dox_NPs, respectively (**Figure 6.F-H**), when compared to PBS-injected embryos. Interestingly, higher HSA coating onto the NP surface increased anti-metastatic effects up to 26% and 49% for NPs and Dox_NPs, respectively, when compared to HSA NPs. In a recent work with bovine serum albumine (BSA) nanoparticles (without a hard, inorganic core, as here employed), Cakan-Akdogan *et al.* showed that these BSA-NPs delivered intravenously to the zebrafish larvae were able to target circulating tumor cells (CTCs) in a zebrafish xenograft model^92^. In addition, the presence of DOX encapsulated inside these BSA-NPs was related to CTC cell death. These observations are in agreement with the colocalization found in this study between Zmel cells and HSA NPs (**Figure S4.B**) and also with the dampening in tumor progression upon treatment with the NPs. As HSA is known to activate the innate immune system through Toll-like receptor 4 (TLR4) and induce inflammation^93–97^, and also as observed in **Figure S3.C**, where approximately 35% of the NPs were internalized by macrophages at 24 hpi, we speculate that HSA-coated NPs may activate a prompt immune response that leads to effective NP internalization and removal from circulation.

Noteworthy, these High HSA NPs enabled a significant gain in survival at 4 dpi in the tumor-grafted zebrafish (with and without DOX, **Figure 6.I**), that puts them close to the control injected with PBS.

Altogether, these results suggest that i) HSA NPs impair metastatic outgrowth independently of the presence of encapsulated DOX and that ii) increasing HSA coverage of NPs improves both, the reduction in tumor volume and the overal survival of embryos. Here, the size and the mere presence of the NPs in circulation, their aggregation at short times in biological media, their strong interactions with the endothelium *in vivo,* and with the Zmel cells *in vitro* and also *in vivo*, might altogether contribute to the anti-tumoral effects observed. Nevertheless, the specific biological pathways and mechanisms behind the here-observed reduction in tumor outgrowth remain to be elucidated.

## Conclusion

Altogether, these results confirm the suitability and biosafety of the HSA-coated IO STMS core-shell NPs as well as their therapeutic efficacy in a syngeneic zebrafish melanoma model. The outgrowth of the tumor mets was markedly halted when the HSA-coated NPs were intravascularly injected in the zebrafish embryos pre-inoculated with melanoma cells. This anti-metastatic effect was evidenced regardless of the encapsulated drug. By increasing the amount of HSA immobilized on the NPs, a significant improvement was observed in the anti-tumoral effect and also in the overall survival of the xenografted animals. The core-shell nanoparticles surface-modified with High HSA appear therefore as promising nanovehicles for drug encapsulation, fertile interactions with tumoral cells, tumor targetting and localized magnetic/NIR hyperthermia treatments.

## Supporting information

All supplemental data

## Acknowledgements

We thank all members of the Tumor Biomechanics Lab for their constant and helpful discussions throughout this investigation. We thank Pascal Kessler (PICSTRA, CRBS) for assistance in imaging and image analysis, and the members of the Mertz lab for their assistance with nanoparticles. D.M. acknowledges the Canceropole Est (project VIVIRMAG) for financial supports. The transmission electronic microscopy platform of the IPCMS is acknowledged for technical support. MT thanks the University of Strasbourg, the National Scientific and Technical Research Council (CONICET), and the Ministry of Science, Technology and Innovation (Mincyt) for financial support. We are particularly grateful to Florent Colin who was pivotal in many parts of this study. Work and people in the Tumor Biomechanics Lab are mostly supported by the INCa (Institut National Du Cancer, French National Cancer Institute), charities (La Ligue contre le Cancer, ARC (Association pour la Recherche contre le Cancer), FRM (Fondation pour la Recherche Médicale)), the National Plan Cancer initiative, the Region Grand Est, INSERM and the University of Strasbourg and from local donators (TDLR, Club Féminin Lampertheim). This work has been directly funded by the support from the Canceropole Grand-Est, with additional support from INCa. VM has been funded by ARC.

## References

1. Pelaz, B. et al. Nano Focus Diverse Applications of Nanomedicine. (2017) doi:10.1021/acsnano.6b06040.

2. Torres Andón, F. & Fadeel, B. Nanotoxicology: Towards Safety by Design. in 391–424 (2014). doi:10.1007/978-3-319-08084-0_14.

3. Sun, T. et al. Engineered nanoparticles for drug delivery in cancer therapy. Angew Chem Int Ed Engl 53, 12320–12364 (2014).

4. Janjua, T. I., Cao, Y., Yu, C. & Popat, A. Clinical translation of silica nanoparticles. Nat Rev Mater 6, 1072–1074 (2021).

5. Mamaeva, V., Sahlgren, C. & Lindén, M. Mesoporous silica nanoparticles in medicine—Recent advances. Adv Drug Deliv Rev 65, 689–702 (2013).

6. Wu, S.-H., Mou, C.-Y. & Lin, H.-P. Synthesis of mesoporous silica nanoparticles. Chem. Soc. Rev. 42, 3862–3875 (2013).

7. Vallet-Regí, M., Schüth, F., Lozano, D., Colilla, M. & Manzano, M. Engineering mesoporous silica nanoparticles for drug delivery: where are we after two decades? Chem. Soc. Rev. 51, 5365–5451 (2022).

8. Croissant, J. G., Fatieiev, Y. & Khashab, N. M. Degradability and Clearance of Silicon, Organosilica, Silsesquioxane, Silica Mixed Oxide, and Mesoporous Silica Nanoparticles. Advanced Materials 29, 1604634 (2017).

9. Adam, A. et al. Orienting the Pore Morphology of Core-Shell Magnetic Mesoporous Silica with the Sol-Gel Temperature. Influence on MRI and Magnetic Hyperthermia Properties. Molecules 26, (2021).

10. Perton, F. et al. Fluorescent and Magnetic Stellate Mesoporous Silica for Bimodal Imaging and Magnetic Hyperthermia. Appl Mater Today 16, 301–314 (2019).

11. Adam, A. et al. Core-shell iron oxide@stellate mesoporous silica for combined near-infrared photothermia and drug delivery: Influence of pH and surface chemistry. Colloids Surf A Physicochem Eng Asp 640, 128407 (2022).

12. Fenaroli, F. et al. Nanoparticles as Drug Delivery System against Tuberculosis in Zebrafish Embryos: Direct Visualization and Treatment. ACS Nano 8, 7014–7026 (2014).

13. Jiang, X.-Y. et al. Quantum dot interactions and flow effects in angiogenic zebrafish (Danio rerio) vessels and human endothelial cells. Nanomedicine 13, 999–1010 (2017).

14. Chakraborty, C., Sharma, A. R., Sharma, G. & Lee, S. S. Zebrafish : A complete animal model to enumerate the nanoparticle toxicity. J Nanobiotechnology 1–13 (2016) doi:10.1186/s12951-016-0217-6.

15. Lin, S., Zhao, Y., Nel, A. E. & Lin, S. Zebrafsh: An In Vivo Model for Nano EHS Studies. Small 9, 1608–1618 (2013).

16. Robin A. Nadar, Nandini Asokan, L. D. E., Alessandra Curci, Alessandra Barbanente, Lukas Schlatt, U. K., Michele Iafisco, d Nicola Margiotta, e Michael Brand, B., Jeroen J. J. P. van den, Beucken, M. B. and & Leeuwenburgh, S. C. G. Preclinical Evaluation of Platinum-Loaded Hydroxyapatite Nanoparticles in an embryonic zebrafish xenograft model Preclinical evaluation of platinum-loaded hydroxyapatite nanoparticles in an embryonic zebra fi sh xenograft model. Nanoscale (2020) doi:10.1039/D0NR04064A.

17. Evensen, L. et al. Zebrafish as a model system for characterization of nanoparticles against cancer. Nanoscale 8, 862–877 (2016).

18. Sieber, S. et al. Zebrafish as an early stage screening tool to study the systemic circulation of nanoparticulate drug delivery systems in vivo. J Control Release 264, 180–191 (2017).

19. Ko, S.-K., Chen, X., Yoon, J. & Shin, I. Zebrafish as a good vertebrate model for molecular imaging using fluorescent probes. Chem. Soc. Rev. 40, 2120–2130 (2011).

20. Evensen, L. et al. Zebrafish as a model system for characterization of nanoparticles against cancer. Nanoscale 8, 862–877 (2016).

21. Dal, N.-J. K. et al. Zebrafish Embryos Allow Prediction of Nanoparticle Circulation Times in Mice and Facilitate Quantification of Nanoparticle–Cell Interactions. Small 16, 1906719 (2020).

22. Johansen, P. L., Fenaroli, F., Evensen, L., Griffiths, G. & Koster, G. Optical micromanipulation of nanoparticles and cells inside living zebrafish. Nat Commun 1–8 (2016) doi:10.1038/ncomms10974.

23. Fenaroli, F. et al. Nanoparticles as Drug Delivery System against Tuberculosis in Zebrafish Embryos: Direct Visualization and Treatment. ACS Nano 8, 7014–7026 (2014).

24. Haldi, M., Ton, C., Seng, W. L. & McGrath, P. Human melanoma cells transplanted into zebrafish proliferate, migrate, produce melanin, form masses and stimulate angiogenesis in zebrafish. Angiogenesis 9, 139–151 (2006).

25. Konantz, M. et al. Zebrafish xenografts as a tool for in vivo studies on human cancer. 1266, 124–137 (2012).

26. Asokan, N. et al. Long-term in vivo imaging reveals tumor-specific dissemination and captures host tumor interaction in zebrafish xenografts. Sci Rep 10, 1–14 (2020).

27. Tavares Barroso, M., et al. Establishment of Pancreatobiliary Cancer Zebrafish Avatars for Chemotherapy Screening. Cells 10, (2021).

28. van Rooijen, E., Fazio, M. & Zon, L. I. From fish bowl to bedside: The power of zebrafish to unravel melanoma pathogenesis and discover new therapeutics. Pigment Cell Melanoma Res 30, 402–412 (2017).

29. Kocere, A. et al. Real-time imaging of polymersome nanoparticles in zebrafish embryos engrafted with melanoma cancer cells: Localization, toxicity and treatment analysis. EBioMedicine 58, 102902 (2020).

30. Tavares Barroso, M., et al. Establishment of Pancreatobiliary Cancer Zebrafish Avatars for Chemotherapy Screening. Cells 10, (2021).

31. Haque, E. & Ward, A. C. Zebrafish as a Model to Evaluate Nanoparticle Toxicity. Nanomaterials (Basel) 8, (2018).

32. Asharani, P. V, Lian Wu, Y., Gong, Z. & Valiyaveettil, S. Toxicity of silver nanoparticles in zebrafish models. Nanotechnology 19, 255102 (2008).

33. Mal, J. et al. A comparison of fate and toxicity of selenite, biogenically, and chemically synthesized selenium nanoparticles to zebrafish (Danio rerio) embryogenesis. Nanotoxicology 11, 87–97 (2017).

34. Bonfanti, P. et al. Functional silver-based nanomaterials affecting zebrafish development: the adverse outcomes in relation to the nanoparticle physical and chemical structure. Environ. Sci.: Nano 11, 2521–2540 (2024).

35. Mutalik, C. et al. Zebrafish Insights into Nanomaterial Toxicity: A Focused Exploration on Metallic, Metal Oxide, Semiconductor, and Mixed-Metal Nanoparticles. Int J Mol Sci 25, (2024).

36. Nadrah, P. et al. Hindered Disulfide Bonds to Regulate Release Rate of Model Drug from Mesoporous Silica. ACS Appl Mater Interfaces 5, 3908–3915 (2013).

37. Velikova, N. et al. Broadening the antibacterial spectrum of histidine kinase autophosphorylation inhibitors via the use of ε-poly-L-lysine capped mesoporous silica-based nanoparticles. Nanomedicine 13, 569–581 (2017).

38. Paatero, I. et al. Analyses in zebrafish embryos reveal that nanotoxicity profiles are dependent on surface-functionalization controlled penetrance of biological membranes. Sci Rep 7, 8423 (2017).

39. Sharif, F., Porta, F., Meijer, A. H., Kros, A. & Richardson, M. K. Mesoporous silica nanoparticles as a compound delivery system in zebrafish embryos. Int J Nanomedicine 7, 1875–1890 (2012).

40. Nasrallah, G. K. et al. A systematic investigation of the bio-toxicity of core-shell magnetic mesoporous silica microspheres using zebrafish model. Microporous and Mesoporous Materials 265, 195–201 (2018).

41. Liu, T.-P., Wu, S.-H., Chen, Y.-P., Chou, C.-M. & Chen, C.-T. Biosafety evaluations of well-dispersed mesoporous silica nanoparticles: towards in vivo-relevant conditions. Nanoscale 7, 6471–6480 (2015).

42. Brown, J. M., Recht, L. & Strober, S. The Promise of Targeting Macrophages in Cancer Therapy. Clin Cancer Res 23, 3241–3250 (2017).

43. Poh, A. R. & Ernst, M. Targeting Macrophages in Cancer: From Bench to Bedside. Front Oncol 8, 49 (2018).

44. Pang, L. et al. Exploiting macrophages as targeted carrier to guide nanoparticles into glioma. Oncotarget 7, 37081–37091 (2016).

45. Patel, S. K. & Janjic, J. M. Macrophage targeted theranostics as personalized nanomedicine strategies for inflammatory diseases. Theranostics 5, 150–172 (2015).

46. Zhao, P. et al. Dual-Targeting to Cancer Cells and M2 Macrophages via Biomimetic Delivery of Mannosylated Albumin Nanoparticles for Drug-Resistant Cancer Therapy. Adv Funct Mater 27, 1700403 (2017).

47. Xie, Z. et al. Targeting tumor hypoxia with stimulus-responsive nanocarriers in overcoming drug resistance and monitoring anticancer efficacy. Acta Biomater 71, 351–362 (2018).

48. Van Driessche, A. et al. pH-Sensitive Hydrazone-Linked Doxorubicin Nanogels via Polymeric-Activated Ester Scaffolds: Synthesis, Assembly, and In Vitro and In Vivo Evaluation in Tumor-Bearing Zebrafish. Chemistry of Materials 30, 8587–8596 (2018).

49. Cakan-Akdogan, G., Ersoz, E., Sozer, S. C. & Gelinci, E. An in vivo zebrafish model reveals circulating tumor cell targeting capacity of serum albumin nanoparticles. J Drug Deliv Sci Technol 75, 103658 (2022).

50. Bozzer, S. et al. Zebrafish: A Useful Animal Model for the Characterization of Drug-Loaded Polymeric NPs. Biomedicines 10, (2022).

51. Kocere, A. et al. Real-time imaging of polymersome nanoparticles in zebrafish embryos engrafted with melanoma cancer cells: Localization, toxicity and treatment analysis. EBioMedicine 58, 102902 (2020).

52. Bizeau, J. et al. Tailoring the pore structure of iron oxide core@stellate mesoporous silica shell nanocomposites: effects on MRI and magnetic hyperthermia properties and applicability to anti-cancer therapies. Nanoscale 16, 15585–15614 (2024).

53. Bizeau, J. et al. Protein sustained release from isobutyramide-grafted stellate mesoporous silica nanoparticles. Int J Pharm X 4, 100130 (2022).

54. Papini, E., Tavano, R. & Mancin, F. Opsonins and Dysopsonins of Nanoparticles: Facts, Concepts, and Methodological Guidelines. Front Immunol 11, 567365 (2020).

55. Owens, D. E. & Peppas, N. A. Opsonization, biodistribution, and pharmacokinetics of polymeric nanoparticles. Int J Pharm 307, 93–102 (2006).

56. Bizeau, J. et al. Protein sustained release from isobutyramide-grafted stellate mesoporous silica nanoparticles. Int J Pharm X 4, 100130 (2022).

57. Chi, N. C. et al. Foxn4 directly regulates. Genes Dev 734–739 (2008) doi:10.1101/gad.1629408.734.

58. Ellett, F., Pase, L., Hayman, J. W., Andrianopoulos, A. & Lieschke, G. J. Mpeg1 Promoter Transgenes Direct Macrophage-Lineage Expression in Zebrafish. Blood 117, e49–e56 (2011).

59. Davison, J. M. et al. Transactivation from Gal4-VP16 transgenic insertions for tissue-specific cell labeling and ablation in zebrafish. Dev Biol 304, 811–824 (2007).

60. Mary, B., Ghoroghi, S., Hyenne, V. & Goetz, J. G. Live tracking of extracellular vesicles in larval zebrafish. in Methods in Enzymology (2020). doi:10.1016/bs.mie.2020.07.013.

61. Hyenne, V. et al. Studying the Fate of Tumor Extracellular Vesicles at High Spatiotemporal Resolution Using the Zebrafish Embryo. Dev Cell 48, 554–572.e7 (2019).

62. Duan, X. & Li, Y. Physicochemical characteristics of nanoparticles affect circulation, biodistribution, cellular internalization, and trafficking. Small 9, 1521–1532 (2013).

63. Blanco, E., Shen, H. & Ferrari, M. Principles of nanoparticle design for overcoming biological barriers to drug delivery. Nat Biotechnol 33, 941–951 (2015).

64. Pensado-López, A. et al. Zebrafish Models for the Safety and Therapeutic Testing of Nanoparticles with a Focus on Macrophages. Nanomaterials (Basel) 11, (2021).

65. Haque, E. & Ward, A. C. Zebrafish as a Model to Evaluate Nanoparticle Toxicity. Nanomaterials (Basel) 8, (2018).

66. Asharani, P. V, Lian Wu, Y., Gong, Z. & Valiyaveettil, S. Toxicity of silver nanoparticles in zebrafish models. Nanotechnology 19, 255102 (2008).

67. Sharif, F., Porta, F., Meijer, A. H., Kros, A. & Richardson, M. K. Mesoporous silica nanoparticles as a compound delivery system in zebrafish embryos. Int J Nanomedicine 7, 1875–1890 (2012).

68. Maeda, H. The enhanced permeability and retention (EPR) effect in tumor vasculature: the key role of tumor-selective macromolecular drug targeting. Adv Enzyme Regul 41, 189–207 (2001).

69. Demirkurt, B., Cakan-Akdogan, G. & Akdogan, Y. Preparation of albumin nanoparticles in water-in-ionic liquid microemulsions. J Mol Liq 295, 111713 (2019).

70. Desai, N. et al. Increased antitumor activity, intratumor paclitaxel concentrations, and endothelial cell transport of cremophor-free, albumin-bound paclitaxel, ABI-007, compared with cremophor-based paclitaxel. Clin Cancer Res 12, 1317–1324 (2006).

71. Ma, P. & Mumper, R. J. Paclitaxel Nano-Delivery Systems: A Comprehensive Review. J Nanomed Nanotechnol 4, 1000164 (2013).

72. Xie, Z. et al. Targeting tumor hypoxia with stimulus-responsive nanocarriers in overcoming drug resistance and monitoring anticancer efficacy. Acta Biomater 71, 351–362 (2018).

73. Cakan-Akdogan, G., Ersoz, E., Sozer, S. C. & Gelinci, E. An in vivo zebrafish model reveals circulating tumor cell targeting capacity of serum albumin nanoparticles. J Drug Deliv Sci Technol 75, 103658 (2022).

74. Tacar, O., Sriamornsak, P. & Dass, C. R. Doxorubicin: an update on anticancer molecular action, toxicity and novel drug delivery systems. J Pharm Pharmacol 65, 157–170 (2013).

75. Cheng, D., Morsch, M., Shami, G. J., Chung, R. S. & Braet, F. Albumin uptake and distribution in the zebrafish liver as observed via correlative imaging. Exp Cell Res 374, 162–171 (2019).

76. Warga, R. M., Kane, D. A. & Ho, R. K. Fate mapping embryonic blood in zebrafish: multi- and unipotential lineages are segregated at gastrulation. Dev Cell 16, 744–755 (2009).

77. Ulanova, L. S. et al. Treatment of Francisella infections via PLGA- and lipid-based nanoparticle delivery of antibiotics in a zebrafish model. Dis Aquat Organ 125, 19–29 (2017).

78. Hyenne, V. et al. Studying the Fate of Tumor Extracellular Vesicles at High Spatiotemporal Resolution Using the Zebrafish Embryo. Dev Cell 48, 554–572.e7 (2019).

79. Tacar, O., Sriamornsak, P. & Dass, C. R. Doxorubicin: an update on anticancer molecular action, toxicity and novel drug delivery systems. J Pharm Pharmacol 65, 157–170 (2013).

80. Cardinale, D., Iacopo, F. & Cipolla, C. M. Cardiotoxicity of Anthracyclines. Front Cardiovasc Med 7, 26 (2020).

81. Etchin, J., Kanki, J. P. & Look, A. T. Zebrafish as a Model for the Study of Human Cancer. Methods Cell Biol 105, 309–337 (2011).

82. Lam, S. H. et al. Conservation of gene expression signatures between zebrafish and human liver tumors and tumor progression. Nat Biotechnol 24, 73–5 (2006).

83. Rothenbücher, T. S. P. et al. Zebrafish embryo as a replacement model for initial biocompatibility studies of biomaterials and drug delivery systems. Acta Biomater 100, 235–243 (2019).

84. Nicoli, S., Ribatti, D., Cotelli, F. & Presta, M. Mammalian Tumor Xenografts Induce Neovascularization in Zebrafish Embryos Mammalian Tumor Xenografts Induce Neovascularization in Zebrafish Embryos. 2927–2931 (2007) doi:10.1158/0008-5472.CAN-06-4268.

85. Van Driessche, A. et al. pH-Sensitive Hydrazone-Linked Doxorubicin Nanogels via Polymeric-Activated Ester Scaffolds: Synthesis, Assembly, and In Vitro and In Vivo Evaluation in Tumor-Bearing Zebrafish. Chemistry of Materials 30, 8587–8596 (2018).

86. Bozzer, S. et al. Zebrafish: A Useful Animal Model for the Characterization of Drug-Loaded Polymeric NPs. Biomedicines 10, (2022).

87. Sambale, F. et al. Three dimensional spheroid cell culture for nanoparticle safety testing. J Biotechnol 205, 120–129 (2015).

88. Peng, Q. et al. Preformed albumin corona, a protective coating for nanoparticles based drug delivery system. Biomaterials 34, 8521–8530 (2013).

89. Bolaños, K., Kogan, M. J. & Araya, E. Capping gold nanoparticles with albumin to improve their biomedical properties. Int J Nanomedicine 14, 6387–6406 (2019).

90. Spada, A., Emami, J., Tuszynski, J. A. & Lavasanifar, A. The Uniqueness of Albumin as a Carrier in Nanodrug Delivery. Mol Pharm 18, 1862–1894 (2021).

91. Elsadek, B. & Kratz, F. Impact of albumin on drug delivery--new applications on the horizon. J Control Release 157, 4–28 (2012).

92. Cakan-Akdogan, G., Ersoz, E., Sozer, S. C. & Gelinci, E. An in vivo zebrafish model reveals circulating tumor cell targeting capacity of serum albumin nanoparticles. J Drug Deliv Sci Technol 75, 103658 (2022).

93. Arroyo, V., García-Martinez, R. & Salvatella, X. Human serum albumin, systemic inflammation, and cirrhosis. J Hepatol 61, 396–407 (2014).

94. David, S. A., Balaram, P. & Mathan, V. I. Characterization of the interaction of lipid A and lipopolysaccharide with human serum albumin: implications for an endotoxin carrier function for albumin. J Endotoxin Res 2, 99–106 (1995).

95. Dziarski, R. Cell-bound albumin is the 70-kDa peptidoglycan-, lipopolysaccharide-, and lipoteichoic acid-binding protein on lymphocytes and macrophages. Journal of Biological Chemistry 269, 20431–20436 (1994).

96. Fukui, H. Relation of endotoxin, endotoxin binding proteins and macrophages to severe alcoholic liver injury and multiple organ failure. Alcohol Clin Exp Res 29, 172S–179S (2005).

97. Jürgens, G. et al. Investigation into the interaction of recombinant human serum albumin with Re-lipopolysaccharide and lipid A. J Endotoxin Res 8, 115–126 (2002).

