## Supplementary material for "Serum albumin coated stellate mesoporous silica nanocomposites inhibit metastatic outgrowth in zebrafish embryos": All supplemental data

1 Tumor Biomechanics Lab, INSERM UMR-S1109, Strasbourg, France.

2 Université de Strasbourg, Strasbourg, France.

3 Fédération de Médecine Translationnelle de Strasbourg (FMTS), Strasbourg, France.

4 Equipe Labellisée Ligue Contre le Cancer, Strasbourg, France.

5 Institut de Physique et Chimie des Matériaux de Strasbourg (IPCMS), UMR-7504 CNRS-Université de Strasbourg, France.

6 Institut de Cancérologie Strasbourg Europe, 67000 Strasbourg, France.

7 Institute of Nanosystems (INS), School of Bio and Nanotechnology, National University of San Martin – CONICET, Buenos Aires, Argentina.

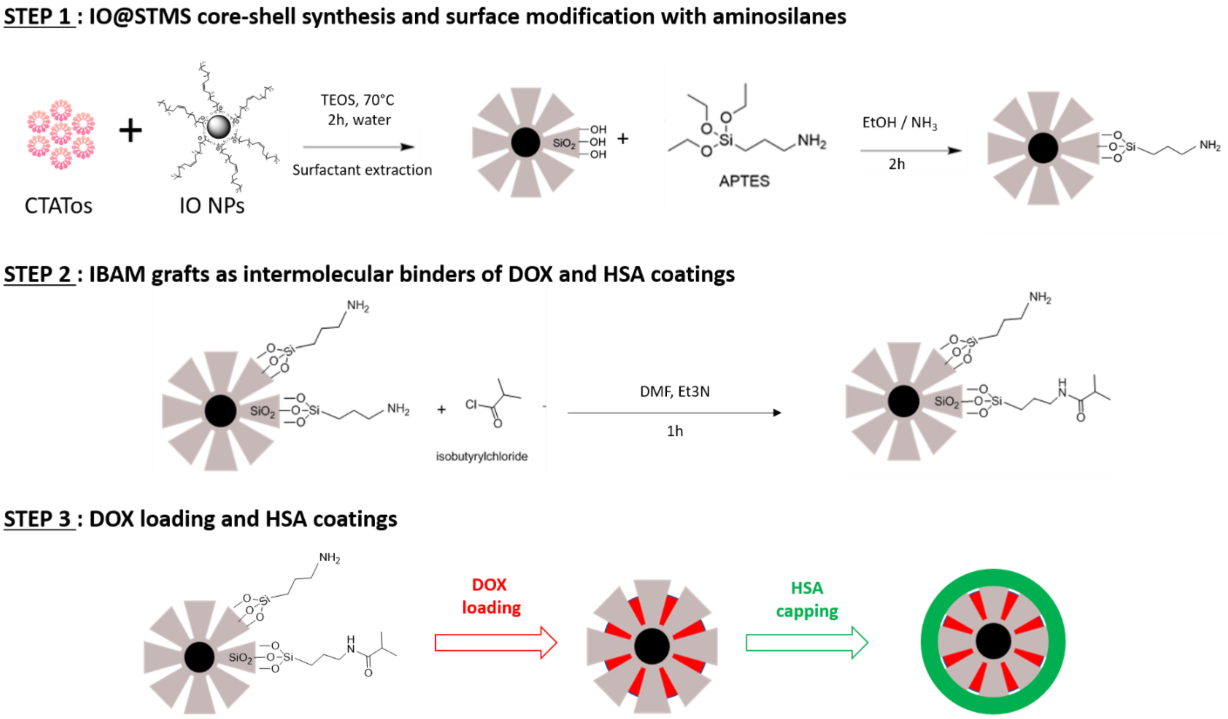


**Scheme S1.** Scheme illustrating the synthesis of the core-shell nanoparticles and the whole surface modification strategy used in this study to load DOX and immobilize HSA.

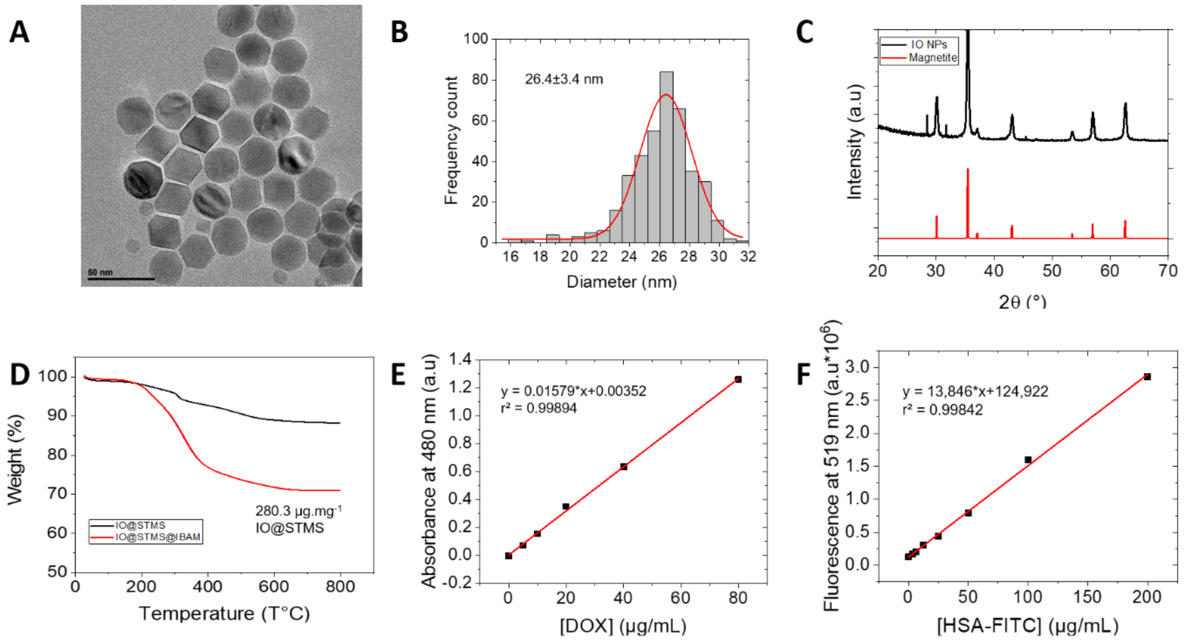


**Figure S1. Physicochemical characterization of the core-shell nanostructures.** A) Transmission electron microscopy image, B) size histogram and C) X-ray diffraction of the iron oxide core nanomaterials. D) Thermogravimetric analysis (TGA) of IBAM-modified IO@STMS NPs (red curve) as compared with IO@STMS NPs (black curve). Standard calibration curves of E) Doxorubicin (480 nm, absorbance) and F) FITC-labelled HSA (519 nm, fluorescence).

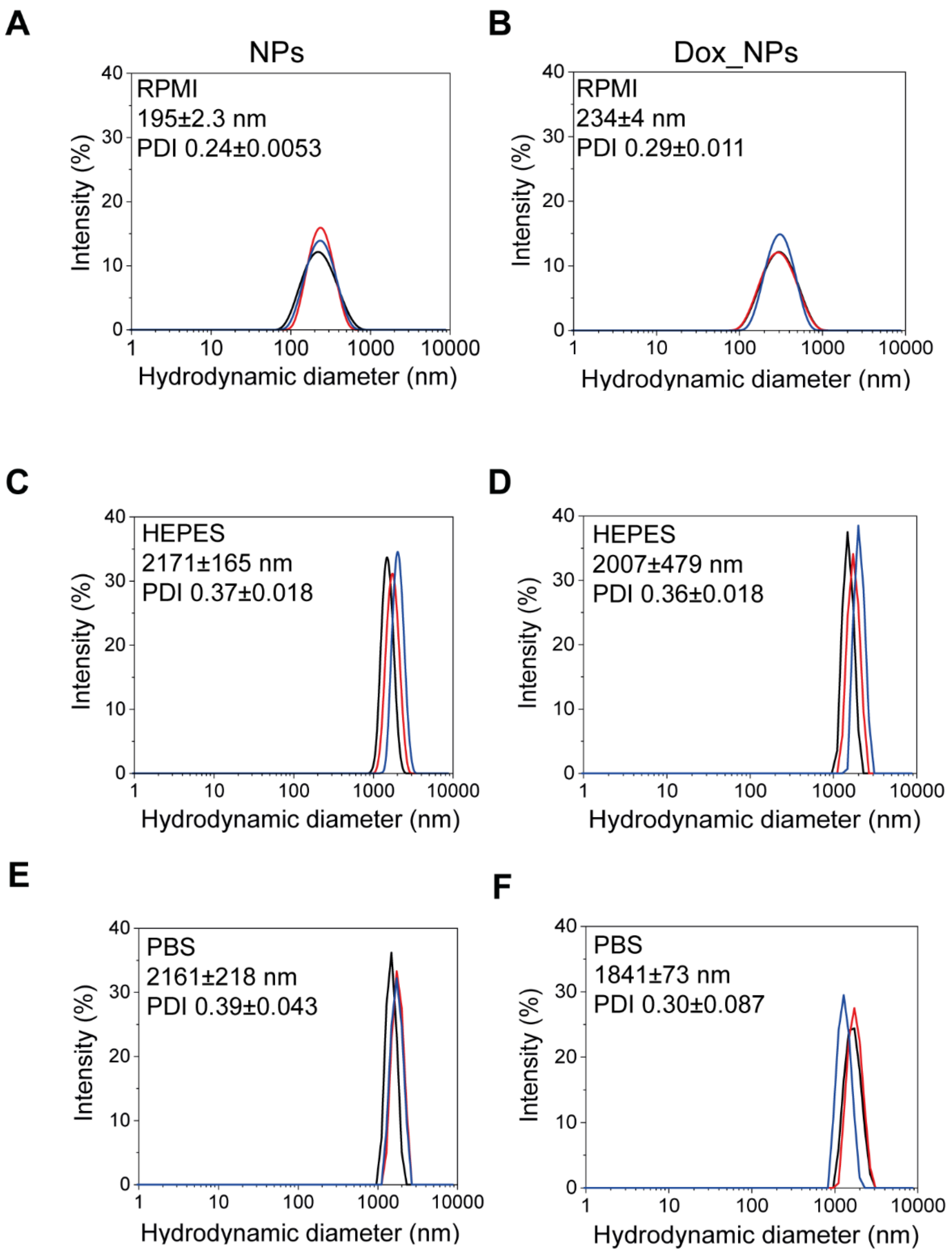


**Figure S2. NP hydrodynamic size distribution in three different biological media for HSA NPs.** DLS intensity measurements in RPMI, HEPES and PBS buffers, respectively, of HSA-FITC NPs (A, C, and E), and HSA-FITC NPs loaded with DOX (B, D, and F). Graphs B-D-F for HSA-FITC Dox_NPs are the same as in Figure 1 and are displayed here to compare with HSA-FITC NPs. DLS measurements were done in triplicate in each medium, as indicated in the DLS graphs.

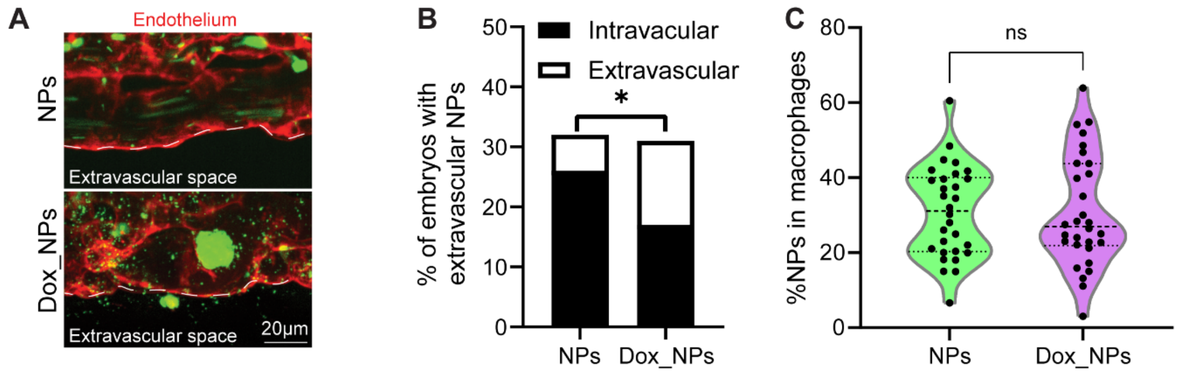


**Figure S3. HSA-Dox NPs are found in the extravascular space in zebrafish embryos.** A) Representative image of the presence of Dox_NPs (bottom) in the extravascular space as delimited by white dotted lines, while their absence is evidenced in the NPs group (top). Tg(Kdrl.hsa-HRAS:mCherry) zebrafish embryos (endothelial cells in red) were injected intravascularly *via* the duct of Cuvier with both NPs and DOX-containing NPs (Dox_NPs) surface-modified with HSA-FITC. B) Quantification of the percentage of embryos showed significant extravascular Dox_NPs when compared to NPs alone (Fisher's exact test, p=0.0319, 3 independent experiments, 30 embryos per group in total). C) Quantification of macrophage internalized NPs showed no difference in colocalization between the two NP groups at 24 hpi (Unpaired t test, p=0,9913, 3 independent experiments, 30 embryos per group in total). Each dot corresponds to the percentage value obtained per each embryo analyzed.

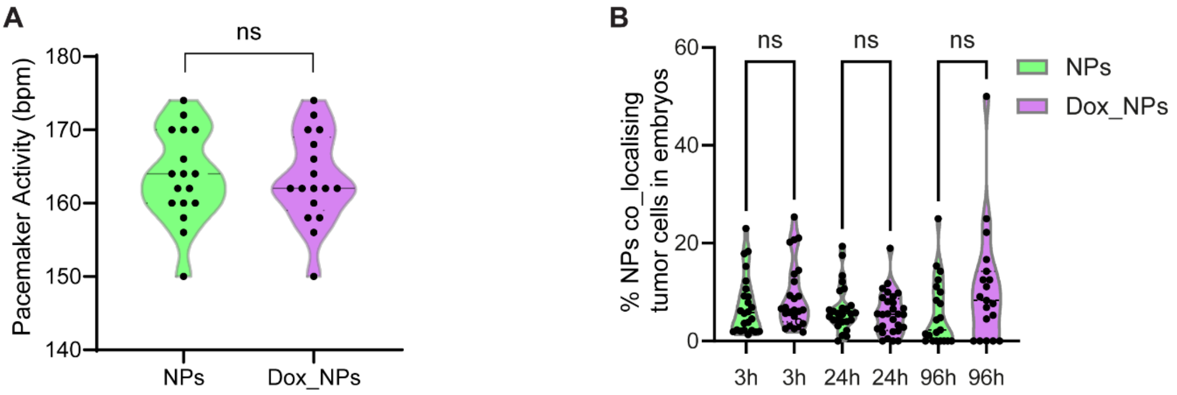


**Figure S4. Quantification of heart beats and of colocalization of NPs with zebrafish melanoma cells *in vivo*.** A) Quantification of heart beats (in beats per minute, bpm) upon microinjected NPs in wildtype zebrafish embryos showed no difference between the two NP groups (Unpaired t test, p=0,8698, 2 independent experiments, 17 embryos per group in total). B) Quantification of NPs colocalizing with zebrafish melanoma cells *in vivo,* showing no difference between NPs and Dox_NPs at the different time points (Mann–Whitney, 3 hpi p=0,2221, 24 hpi p=0,4781, and 96 hpi p=0,1692, 3 independent experiments, minimum 19 cells analyzed per group per time point.

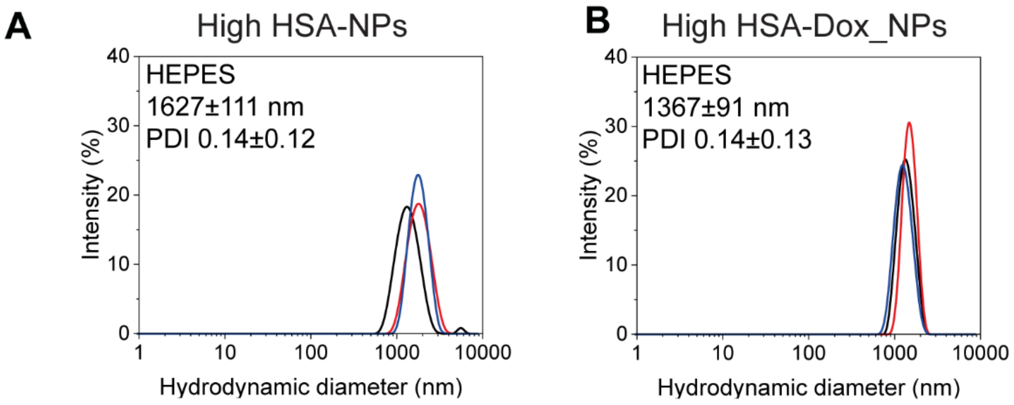


**Figure S5. NP hydrodynamic size distribution in HEPEs for High HSA NPs.** DLS intensity measurements in HEPES buffer for High HSA-FITC NPs (A) and High HSA-FITC-Dox NPs (B). DLS measurements were done in triplicate, as indicated in the DLS graphs.
